# Spatiotemporal biosensor profiling reveals an autonomous mitochondrial NAD^+^/NMN regulatory network centered on NMNAT3

**DOI:** 10.64898/2026.07.11.737903

**Authors:** Liuqing Chen, Xu Cao, Yaqi Wang, Qiuliyang Yu

**Author notes:** Correspondence: Liuqing Chen, Qiuliyang Yu.

## Abstract

Nicotinamide adenine dinucleotide (NAD^+^) and its precursor nicotinamide mononucleotide (NMN) are strictly compartmentalized, yet how individual organelles maintain local metabolic homeostasis remains unresolved. Here, we report FrNADS and FrNMNS^1.0^, a FRET-based biosensor toolkit that maps NAD^+^ and NMN dynamics in living cells with subcellular resolution, including the oxidizing lumen of the endoplasmic reticulum. We find that NAD^+^ recovery in the nucleus following PARP1 activation depends on NAMPT mediated salvage synthesis, while peroxisomes buffer NAD^+^ via NUDT12 and SLC25A17. In mitochondria, NMNAT3 acts as a NAD^+^ hydrolase that counterbalances import through SLC25A51; HINT2 functionally enhances this activity. Furthermore, SLC25A48 functions as a critical regulatory node that modulates the compartmental redistribution of the generated NMN. These findings establish a mitochondrial NAD^+^/NMN regulatory circuit and reveal how organelles independently resolve metabolic stress.

## INTRODUCTION

Nicotinamide adenine dinucleotide (NAD^+^) and its reduced form NADH, together with their phosphorylated counterparts NADP(H), are essential cofactors that govern a wide range of biological processes across all kingdoms of life. NAD^+^ also serves as a substrate for PARPs, CD38, sirtuins, and SARM1^1–4^, linking it to signaling and immunity. Declining NAD^+^ levels mark aging and age-related disease^5,6^, driving interest in NAD^+^-boosting therapies. Understanding of NAD^+^ biology is therefore essential for developing effective interventions.

NAD^+^ is distributed through the cytosol, nucleus, mitochondria, peroxisomes, endoplasmic reticulum (ER), and Golgi apparatus (GA)^7,8^. While NAD^+^ dynamics have been characterized in the cytosol, nucleus and mitochondria, regulation in peroxisomes, the ER and Golgi remain poorly understood. The biosynthesis, localization, and transport of NAD^+^ and its intermediates such as nicotinamide mononucleotide (NMN) are critical for cellular function^9–11^. How compartmentalized NAD^+^ homeostasis influences physiology during aging and stress remains largely unknown^12^.

Recent work has clarified organelle specific NAD^+^ metabolism. For instance, SELO hydrolyzes NAD^+^ to NMN and AMP to modulate lipid metabolism and support mitochondrial homeostasis^13^. This finding suggests that NMN is present within mitochondria. Yet NMN does not effectively raise NAD^+^ in isolated mitochondria^14^. Instead, NAD^+^ enters mitochondria directly through SLC25A51^15–17^, the principal mitochondrial NAD^+^ transporter. These observations argue against a major role for NMN as a precursor for mitochondrial NAD^+^ synthesis.

The physiological role of NMNAT3 has remained equally controversial. This enzyme was traditionally viewed as the mitochondrial NAD^+^ synthase, yet *Nmnat3* knockout mice retain normal mitochondrial NAD^+^ in most tissues^18^. NMNAT3 overexpression elevates mitochondrial NAD^+^ but produce aberrant NAD^+^ analogs, suggesting a buffering or chaperone function rather than net synthesis^19^. A recent model proposes that subcellular NAD^+^ pools are interconnected and that mitochondria buffer fluctuation by importing NAD^+^ via SLC25A51 while reversibly cleaving NAD^+^ through NMNAT3^8^. This model was inferred from steady-state metabolomics in bulk lysates. Whether NMNAT3 synthesizes or hydrolyzes NAD^+^ in living cells, and how this activity is regulated, has never been observed directly in its native compartment. The NAD^+^/NMN ratio serves as a critical activator of SARM1^20^, a regulator of axonal degeneration and immune signaling. Understanding how mitochondria control this ratio is therefore essential for deciphering compartmentalized NAD^+^ metabolism. Collectively, these observations raise a fundamental question: what is the metabolic fate of NMN generated within mitochondria?

Localized NAD^+^ pools exist in multiple organelles, yet current models often treat organelles as dependent compartments that exchange metabolites during stress. Because bulk NAD^+^ cannot cross membranes and the lack of robust compartment specific sensors, a key question remains unanswered: do organelles possess autonomous buffering and clearance mechanisms, or do they rely primarily on transport between compartments? When NAD^+^ is consumed locally, how do impermeable organelles (such as mitochondria and peroxisome) handle the resulting NMN surge without triggering osmotic or metabolic toxicity?

Genetically encoded biosensors enable metabolite tracking in live cells with subcellular resolution. Existing NAD^+^ sensors—including PARAPLAY^7^, LigA-cpVenus^21^, NAD-Snifit^22^, FiNad^23^, NS-X^24^ and ChemoX-NAD^+25^—have revealed compartmentalized NAD^+^ distribution, cell type specific regulation, and age-dependent decline. However, these sensors suffer from pH sensitivity and cross-reactivity with analogs, limiting their use in peroxisomes, the ER, and the GA. To address this, we developed FrNADS and FrNMNS^1.0^, a FRET-based sensor toolkit that overcomes the ATP sensitivity, divalent cation interference, and oxidizing compartment incompatibility of prior tools. Using this toolkit, we directly visualized NMNAT3 mediated NAD^+^ hydrolysis *in situ*, identified HINT2 as a functional enhancer of NMNAT3 hydrolytic activity, and uncovered SLC25A48 as a regulator of mitochondrial NMN compartmental redistribution. These findings establish a mitochondrial NAD^+^/NMN ratio regulatory circuit and reveal how organelles autonomously resolve metabolic stress.

## Results

### Development and characterization of FrNADS and FrNMNS^1.0^

We engineered FRET-based sensors to monitor NAD^+^ and NMN dynamics in living cells. The NAD^+^ sensor sandwiches the DNA ligase domain from *Enterococcus faecalis* V583 (*Ef*LigA) between mScarlet-I3 and StayGold-E138D (Fig. 1A). Ligand binding changes the conformation of *Ef*LigA. This alters the FRET efficiency and produces an emission ratio (595/520 nm, or RFP/GFP) that reports the analyte concentration. Starting from the scaffold of our previously reported NAD^+^ sensor NS-Goji1.0^24^, we optimized the linker between *Ef*LigA and the fluorescent proteins. The resulting variant, FrNADS^0.1^, showed a 3.1-fold dynamic range (Fig. S1A, 1B). However, it remained sensitive to ATP, a limitation shared by most existing NAD^+^ sensors_21,23–25._

**Figure 1.**
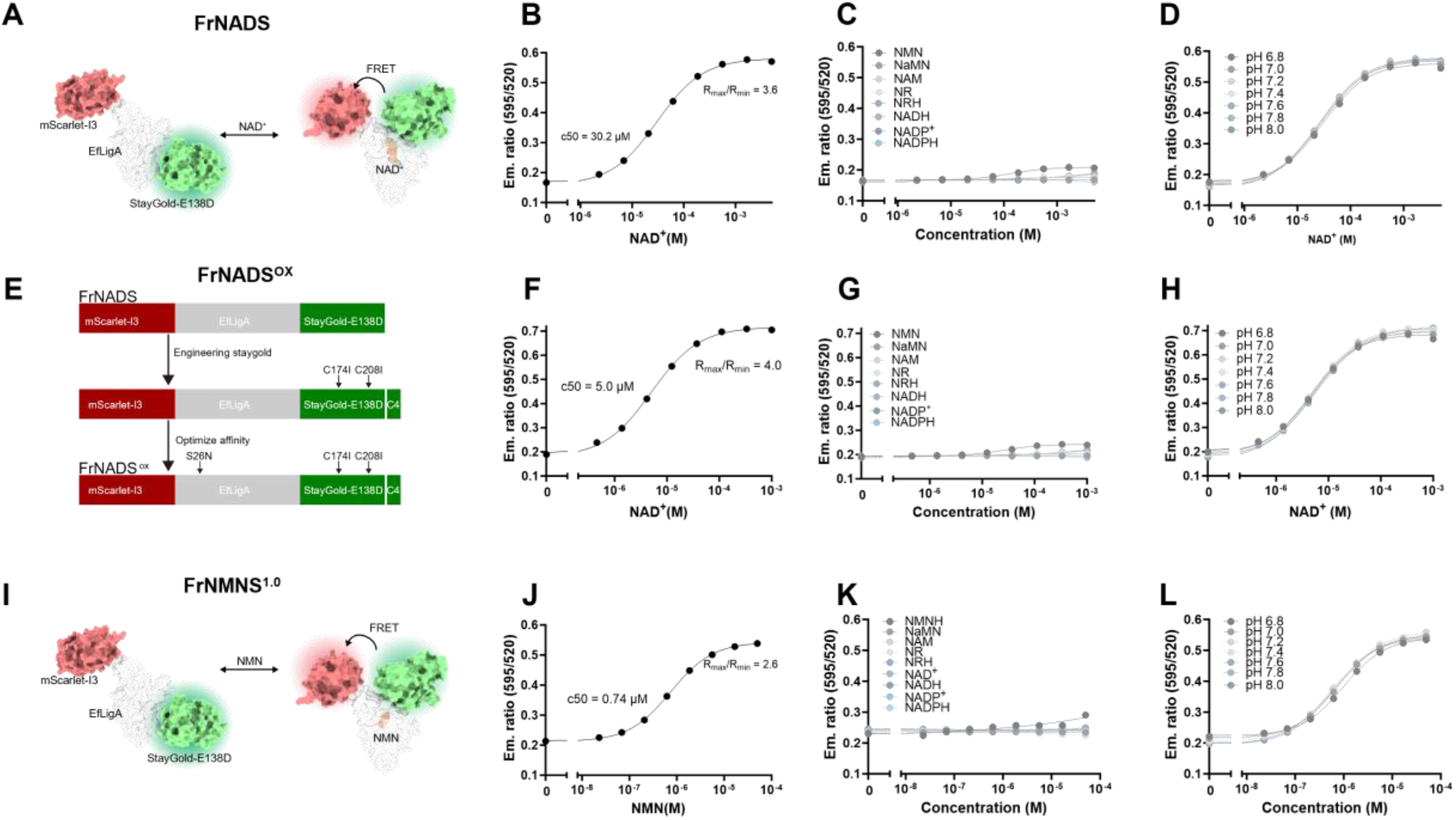
Characterization of biosensor FrNADS and FrNMNS^1.0^. (A) Schematic representation of the FrNADS structure, modeled with the EfLigA domain sandwiched between StayGold-E138D and mScarlet-I3. (B) In vitro dose-response curve of purified FrNADS to NAD^+^ at room temperature (RT), pH 7.2. The sensor exhibits a 3.6-fold emission ratio change with an apparent affinity (c50) of 30.2 μM. (C) FrNADS shows minimal cross reactivity to NAD^+^ analogs (NAM, NR, NRH, NMN, NaMN, NADH, NADP^+^, NADPH). (D) The emission ratio of FrNADS remains stable across the physiological pH range (6.8-8.0). (E) Schematic workflow of the development process of FrNADS^ox^. (F) In vitro dose-response curve of FrNADS^ox^ to NAD^+^ at RT, pH 7.2, with a c50 of 5.0 μM. (G) FrNADS^ox^ shows high specificity. (H) The emission ratio of FrNADS^ox^ remains stable across the physiological pH range (6.8-8.0). (I) Schematic representation of FrNMNS^1.0^ structure. (J) In vitro dose-response curve of purified FrNMNS^1.0^ to NMN at RT, pH 7.2, with a c50 of 0.74 μM. (K) FrNMNS^1.0^ does not respond to NMN analogs (NAM, NR, NRH, NMNH, NaMN, NAD^+^, NADH, NADP^+^, NADPH). (L) The emission ratio of FrNMNS^1.0^ is insensitive to pH changes within the physiological range (6.8-8.0). Data are presented as mean ± SD from at least three independent measurements.

Existing NAD^+^ sensors have not been systematically tested against divalent cations^21,23–25^. Mg^2+^ and Ca^2+^ markedly shifted the apparent NAD^+^ affinity of FrNADS^0.1^ (Fig. S1C, S1D). We reasoned that charged residues near the binding pocket mediate this effect. Mutating Asp91, a negatively charged residue adjacent to the binding site, produced variants with altered cation sensitivity (Fig. S1E). The D91Q mutation achieved the best balance. It reduced cation interference substantially while retaining high selectivity (Fig. S1D-I). In contrast, D91L eliminated cation sensitivity but caused unacceptable cross-reactivity with NMN. The final NAD^+^ sensor, FrNADS, carries an additional R202A mutation to further reduce ATP interference.

Purified FrNADS showed a 3.6-fold emission ratio change in response to NAD^+^, with an apparent half maximal effective concentration (c50) of 30.2 μM. It discriminated well against NAD^+^ analogs and showed minimal pH sensitivity across the physiological range (Fig. 1B, 1C, S2A). Mg^2+^ and Ca^2+^ did not affect its response (Fig. S2B). Adenosine nucleotides (AXPs) caused minor interference (Fig. S2C), but the total cellular AXP pool is relatively stable under physiological conditions. Furthermore, changes in the ADP/ATP ratio did not alter FrNADS performance (Fig. S2D). The emission ratio was independent of sensor concentration (Fig. S2E), which is essential for reliable intracellular measurements. Temperature had a modest effect on sensor performance (Fig. S2F).

To extend the sensor toolkit to oxidizing compartments such as the ER and GA, we introduced additional mutations (C174I, C208I, and S26N). These changes improve oxygen resistance and tuned ligand affinity (Fig. 1E). The resulting variant, FrNADS^ox^, bound NAD^+^ with a c50 of 5.0 μM. It retained high selectivity and pH tolerance (Fig. S1F-H, S2J), and showed minimal temperature sensitivity (Fig. S2K). AXPs modestly affected its NAD^+^ affinity (Fig. S2H), but the stable total AXP pool in cells ensures reliable response. The ADP/ATP ratio did not influence its performance (Fig. S2I). Like FrNADS, FrNADS^ox^ showed an emission ratio that was independent of sensor concentration (Fig. S2J).

To monitor multiple nodes in NAD^+^ metabolism, we developed a FRET-based NMN sensor, FrNMNS^1.0^ (Fig. 1I). We used the NMN-binding domain from our previously reported BRET-based NMN sensor^26^. FrNMNS^1.0^ bound NMN with a c50 of 0.74 μM (Fig. 1J). It showed excellent selectivity and pH tolerance (Fig. 1K, 1L, S2L). Cations did not affect its performance (Fig. S2M). Unlike FrNADS, FrNMNS^1.0^ showed minimal interference from AXPs (Fig. S2N), likely due to the absence of the adenosine moiety in NMN. The sensor response was independent of its expression level (Fig. S2O). Temperature caused only minor interference (Fig. S2P). Together, these sensors advance the ability to dissect NAD^+^ biology at subcellular resolution, including in ion-rich or oxidizing microenvironments that existing tools cannot access.

### Spatiotemporal mapping of subcellular NAD^+^ and NMN dynamics

To map subcellular NAD^+^ homeostasis, we targeted FrNADS to six compartments in HEK 293T cells: cytoplasm (Cyto), nucleus (Nuc), mitochondria (Mito), peroxisome (Pex), ER, and GA. After 24 h of treatment with either FK866 (a NAMPT inhibitor) or NR (an NAD^+^ precursor), ratiometric imaging (RFP/GFP) showed that FrNADS responded specifically to NAD^+^ depletion or elevation in every compartment tested (Fig. 2A; representative images in Fig. S3). These results confirm that FrNADS reliably report localized intracellular NAD^+^ pools.

**Figure 2.**
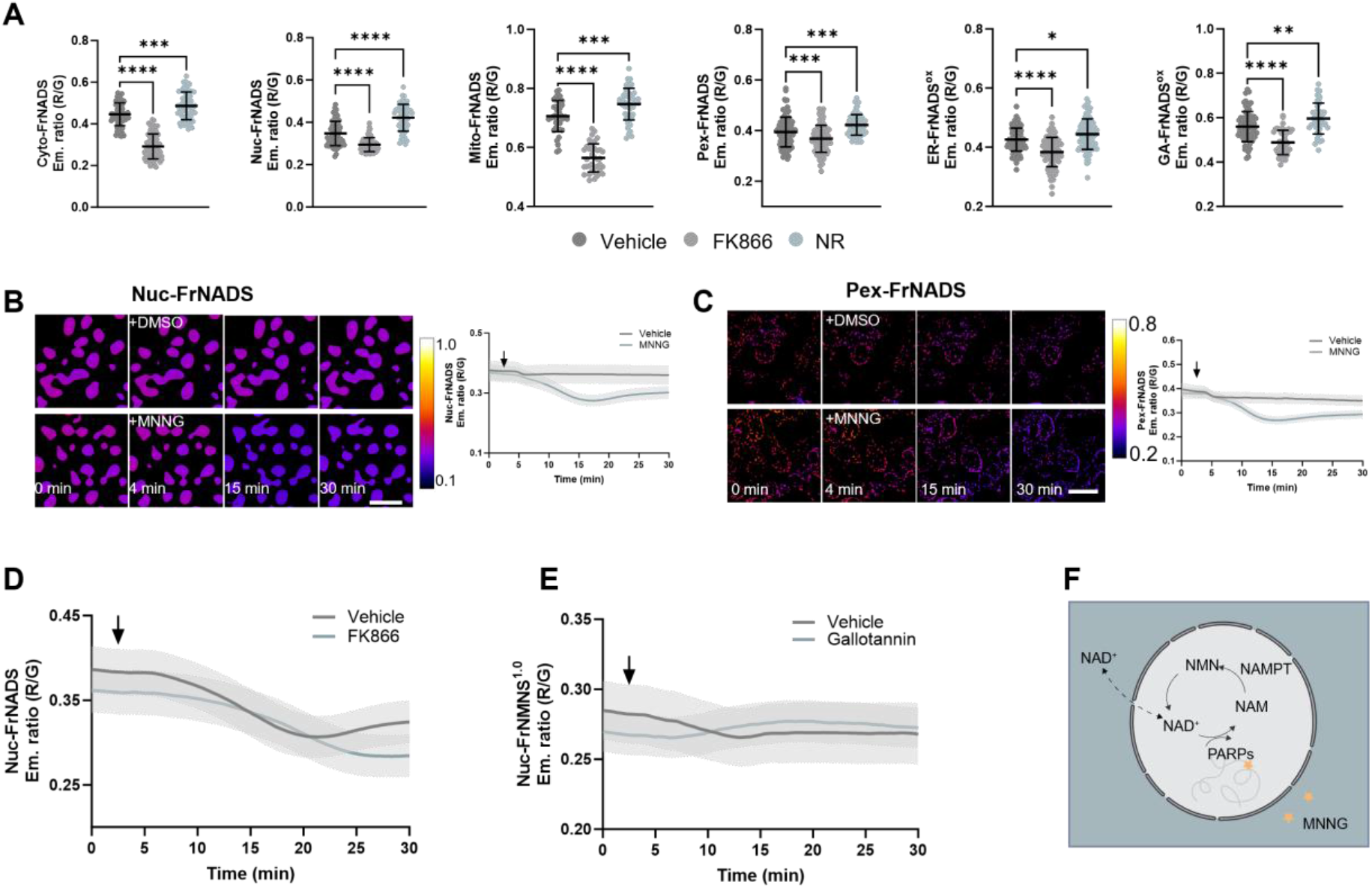
Subcellular NAD^+^ profiling and autonomous replenishment after acute DNA damage. (A) Functional validation of compartmentalized FrNADS biosensors. HEK 293T cells expressing FrNADS targeted to the cytoplasm (Cyto), nucleus (Nuc), mitochondria (Mito), peroxisomes (Pex), endoplasmic reticulum (ER), and Golgi apparatus (GA) were treated with Vehicle, 10 nM FK866 (NAMPT inhibitor), or 500 μM NR (NAD^+^ precursor) for 24 h. The emission ratio (R/G) indicates relative NAD^+^ levels, demonstrating the robust responsiveness of the biosensors across all investigated organelles. (B-C) Real-time tracking of acute subcellular NAD^+^ depletion induced by DNA damage. Representative time-lapse pseudo color images (left panels) and corresponding quantitative kinetic traces (right panels) of FrNADS in nuclear (B) and peroxisome (C). HEK 293T cells were treated with 100 μM MNNG to trigger PARP1 hyperactivation, or with DMSO as a vehicle control. Black arrows indicate the exact time of localized compound addition. Scale bars, 30 μm. (D) Kinetic traces of Nuc-FrNADS in HEK 293T cells treated with MNNG alone or pre-treated (1 h before imaging) with 10 nM FK866 followed by MNNG challenge. Sustained inhibition of NAMPT by FK866 completely abolishes the compensatory NAD^+^ recovery in the nucleus. (E) Kinetic traces of Nuc-FrNMNS^1.0^ in HEK 293T cells treated with MNNG alone or pre-treated (1 h before imaging) with 100 μM gallotannin. Gallotannin inhibit NMNATs, thus enable detect the increase signal of NMN after MNNG challenge. (F) Schematic representation of proposed nuclear NAD^+^ replenish model after acute damage. Data are presented as mean ± SD; statistical significance was determined using one-way ANOVA analysis followed by Dunnett’s multiple comparisons test. ns, P ≥ 0.05; *, P < 0.05; **, P < 0.01; ***, P < 0.001; ****, P < 0.0001.

We next used FrNADS to track acute NAD^+^ fluctuations. We treated cells with MNNG, an alkylating agent that activates PARP1 and triggers rapid NAD^+^ consumption. Time-lapse imaging revealed a rapid decline in NAD^+^ across all compartments monitored (Fig. 2B, 2C, S4A-D). In the nucleus, NAD^+^ reached a minimum approximately 15 min after treatment and then underwent a reproducible partial recovery (Fig. 2B). Peroxisome showed a similar recovery pattern (Fig. 2C). In contrast, NAD^+^ depletion in the other compartments remained sustained for the duration of the experiment.

We hypothesized that the salvage pathway drives this localized NAD^+^ restoration. PARP-mediated NAD^+^ consumption generates nicotinamide (NAM). NAMPT recycles NAM to NMN, and NMNAT converts NMN back to NAD^+^. To test this model, we pre-treated cells with FK866 before adding MNNG. This abolished the partial nucleus recovery completely (Fig. 2D), confirming that the recovery phase requires the NAM salvage pathway. To test the downstream NMN to NAD^+^ conversion step, we expressed FrNMNS^1.0^ in the nucleus and treated cells with NMNAT inhibitor gallotannin before MNNG challenge. We observed a small increase in NMN signal upon gallotannin pretreatment (Fig. 2E), consistent with blocked NMNAT activity. Although gallotannin is a broad-spectrum inhibitor of the NMNAT family and may exert off-target effects, this pharmacological result supports the notion that salvage pathway activity confers the nuclear NAD^+^ recovery. Figure 2F summarizes this model. These dynamics indicate that the nucleus can regenerate NAD^+^ rapidly through local salvage synthesis under our experimental conditions. We cannot exclude a minor contribution from mitochondrial NAD^+^ export^8,27^. However, the complete abolition of recovery by FK866 showed that salvage synthesis is necessary and sufficient to explain the observed replenishment kinetics.

In parallel, we characterized the subcellular performance of FrNMNS^1.0^. Targeted expression in the cytoplasm, nucleus, and mitochondria confirmed that FrNMNS^1.0^ effectively report NMN fluctuations in these distinct compartments (Fig. S5).

Collectively, these findings demonstrate that FrNADS and FrNMNS^1.0^ perform reliably at the subcellular level. The sensitivity of FrNADS enables detection of subtle, compartment-specific NAD^+^ dynamics. This includes the transient, salvage-mediated NAD^+^ recovery in the nucleus after acute DNA damage.

### NAD^+^ and NMN pool regulation at peroxisome

HepG2 cells contain abundant peroxisomes. We used them to study peroxisomal NAD^+^ homeostasis. We targeted FrNADS to peroxisomes and knocked down PXMP2, NUDT12, and the candidate peroxisomal NAD^+^ transporter SLC25A17^28^. Ratiometric imaging showed that NUDT12 knockdown raised steady-state peroxisomal NAD^+^, whereas SLC25A17 knockdown lowered it (Fig. 3A, S6). PXMP2 knockdown had no effect. Overexpression produced the opposite results: NUDT12 reduced peroxisomal NAD^+^, SLC25A17 increased it, and PXMP2 again had no effect (Fig. 3B). These data identify SLC25A17 as a positive regulator of peroxisomal NAD^+^ and NUDT12 as a negative regulator. NUDT12 likely consumes NAD^+^ through its hydrolase activity.

**Figure 3.**
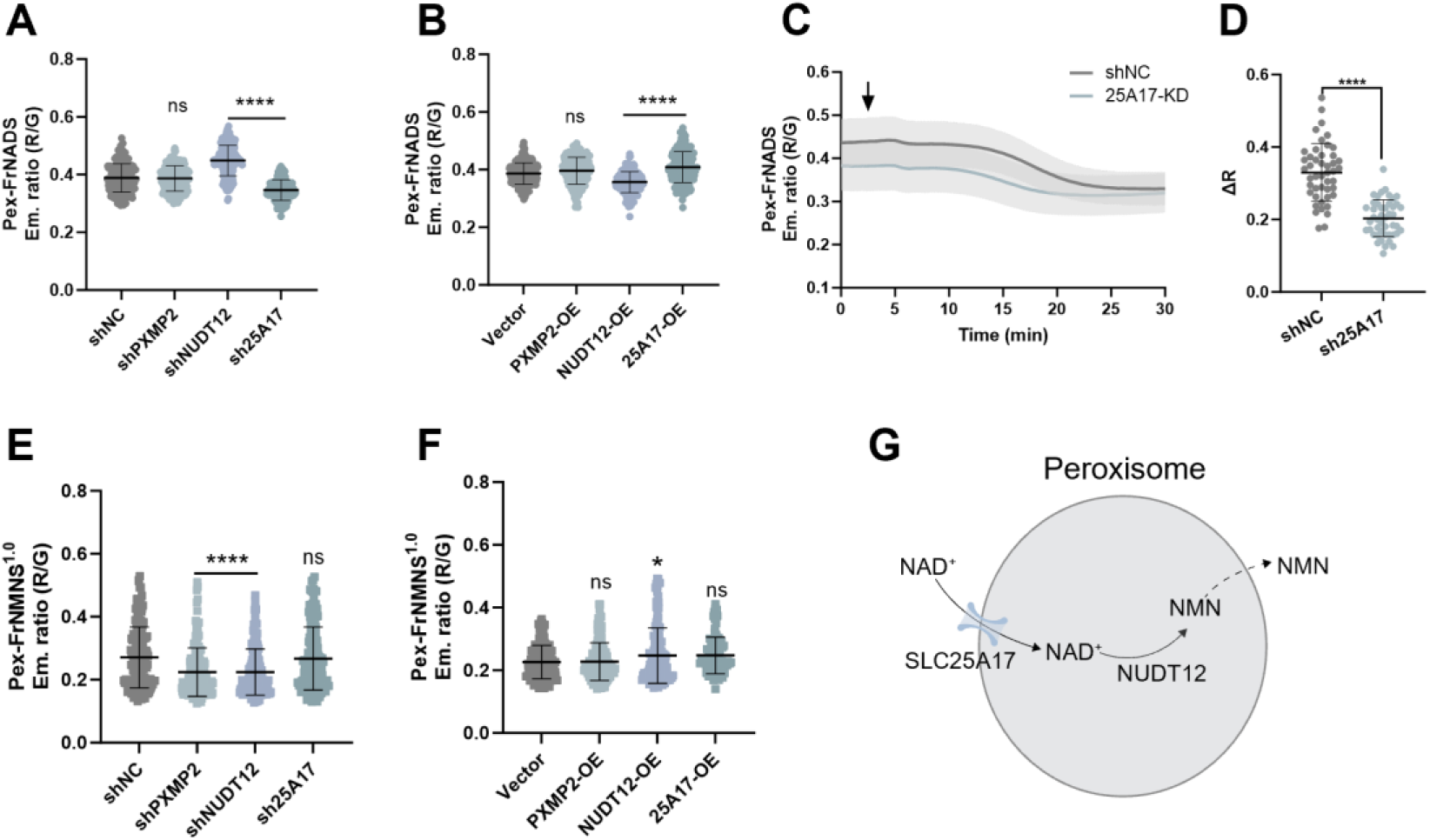
NAD^+^ and NMN regulation at peroxisome. (A and B) Ratiometric quantification of steady-state peroxisome NAD^+^ levels under genetic perturbations. HepG2 cells expressing FrNADS biosensor were subjected to shRNA-mediated knockdown (A) or overexpression (B) of peroxisome candidates (PXMP2, NUDT12, and SLC25A17). (C) continuous real-time time-lapse tracking of Pex-FrNADS in shNC and SLC25A17 knockdown cells. The black arrow indicates the time of MNNG addition. (D) Quantification of the maximum dynamic response amplitude changes (ΔR = R_basal_/R_min_ - 1) extracted from the kinetic traces in (C). (E and F) Reciprocal assessment of interconnected peroxisomal NMN pool utilizing the targeted FrNMNS^1.0^ biosensor. Steady-state emission ratios of FrNMNS^1.0^ in cells with knockdown (E) or overexpression (F) of the indicated components. (G) Schematic representation of proposed peroxisomal NAD^+^ homeostasis regulation mechanism. Data are presented as mean ± SD; statistical significance was determined using one-way ANOVA analysis followed by Dunnett’s multiple comparisons test. ns, P ≥ 0.05; *, P < 0.05; ****, P < 0.0001.

To validate SLC25A17, we monitored NAD^+^ dynamics in real-time. SLC25A17 knockdown cells showed a blunted response to NAD^+^ depletion triggered by MNNG (Fig. 3C, D). NAD^+^ dropped faster and reached baseline sooner than in control cells. This accelerated exhaustion, together with reduced amplitude, points to a smaller basal NAD^+^ pool with limited buffering capacity. SLC25A17 therefore maintains both the steady-state level and the dynamic responsiveness of peroxisomal NAD^+^.

We next examined NMN pool using peroxisomal FrNMNS^1.0^. Knockdown of PXMP2 and NUDT12 both reduced peroxisomal NMN (Fig. 3E). Overexpression of NUDT12 increased NMN (Fig. 3F). Thus, NUDT12 converts NAD^+^ to NMN inside peroxisome. Together with SLC25A17, it governs local NMN and NAD^+^ homeostasis (Fig. 3G).

### Direct mapping of NMNAT3 mediated NAD^+^ hydrolysis in mitochondria

Recent studies proposed that NMNAT3 hydrolyzes NAD^+^ in mitochondria^8,13,29^. That conclusion rested on steady state metabolomics of cell lysates or indirect changes in bulk NAD^+^. Equipped with the FrNMNS^1.0^ and FrNADS sensors, we directly examined NMNAT3 catalytic activity *in situ*.

We first made a mutant, R206A, that lost NAD^+^ hydrolysis activity but retained residual synthetic activity *in vitro* (Fig. 4A and B). We confirmed NMNAT3 knockdown efficiency in purified HepG2 mitochondria and verified the expression of WT and R206A mutants by immunoblotting (Fig. 4C, D). Knockdown significantly reduced matrix NMN and raised NAD^+^. Re-expression of WT NMNAT3 restored both metabolites, but R206A did not (Fig. 4E and F). We further corroborated these sensor-based readouts by LC-MS analysis of immunopurified mitochondria (Fig. S7). We also confirmed NMN accumulation dependent on NMNAT3 in NMNAT3 knockout HEK 293T cells (Fig. S8 and S9A). Notably, NMNAT3 loss did not change mitochondrial NAD^+^ in HEK 293T (Fig. S9B), which is consistent with previous reports^8,18^, and likely reflects the low expression of NMNAT3 in this cell line. While endogenous NMNAT3 is detectable by western blotting in HepG2.

**Figure 4.**
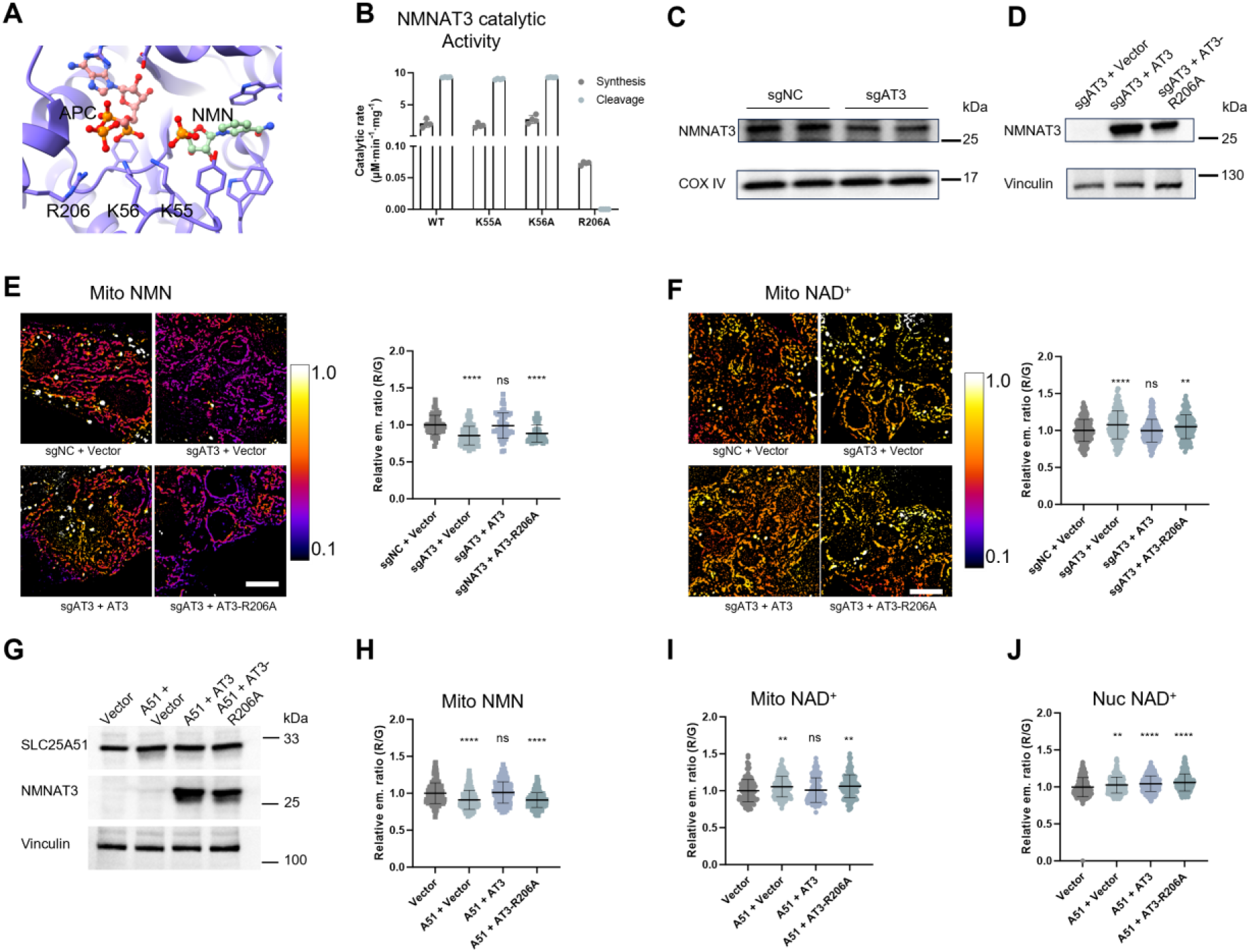
NMNAT3 function as NAD^+^ hydrolyzing enzyme in HepG2 mitochondria. (A) Molecular structural model of NMNAT3 in complex with APC and NMN (PDB code: 1nus), illustrating the substrate binding pocket and interacted residues. (B) NMNAT3 R206A mutation abolished hydrolytic activity, while remain weak synthetic activity. (C and D) Immunoblotting analysis of knockdown and overexpression efficiency of NMNAT3 in HepG2 cells using purified mitochondria. (E and F) Representative pseudocolor FRET emission ratio (RFP/GFP) images of HepG2 cells expressing mitochondrial matrix targeting FrNMNS^1.0^ (E) or FrNADS (F), illustrating the relative levels of mitochondrial NMN and NAD^+^ under NMNAT3 knockdown (sgAT3) or overexpression conditions, compared to non-targeting control (sgNC). Scale bars, 2.5 μm. (G) Immunoblotting analysis of overexpression efficiencies of SLC25A51 and NMNAT3 in HepG2. (H, I, and J) Quantitative analysis of relative mitochondrial NMN (H), mitochondrial NAD^+^ (I), and Nucleus NAD^+^ (J) levels in HepG2 overexpressing of SLC25A51 and NMNAT3. Data are presented as mean ± SD; statistical significance was determined using one-way ANOVA analysis followed by Dunnett’s multiple comparisons test or two-tailed Student’s t-test. ns, P ≥ 0.05; **, P < 0.01; ****, P < 0.0001.

SLC25A51 and NMNAT3 may cooperate to balance mitochondrial NAD^+8,13^. To test this, we overexpressed SLC25A51 in HepG2 cells (Fig. 4G), which decreased matrix NMN and increased NAD^+^ as expected. Co-expression of WT NMNAT3, but not R206A, reversed these changes (Fig. 4H, I). This epistasis shows that NMNAT3 hydrolysis can counteract SLC25A51 import. It acts as a tunable clearance mechanism in the matrix.

We next asked whether this mitochondrial hydrolysis influences nuclear NAD^+^ pools. Unexpectedly, NMNAT3 failed to reverse the nuclear NAD^+^ elevation caused by SLC25A51 overexpression (Fig. 4J). This supports the autonomy of mitochondrial NAD^+^ regulation. In summary, NMNAT3 is a bona fide NAD^+^ hydrolase in the mitochondrial matrix. It offset SLC25A51-dependent import to prevent hyperaccumulation and preserve local NAD^+^/NMN stoichiometry.

### NMNAT3 modulates mitochondrial metabolism via localized NAD^+^ buffering

Given NMNAT3’s primary role as a mitochondrial NAD^+^ hydrolase, we investigated its impact on metabolic homeostasis. NMNAT3 knockdown promoted mitochondrial fatty acid oxidation (FAO) activity (Fig. 5A) and reduced lipid accumulation in HepG2 cells, as shown by Oil Red O staining (Fig. 5B, C). Thus, NMNAT3 mediated NAD^+^ clearance modulates mitochondrial lipid catabolism.

**Figure 5.**
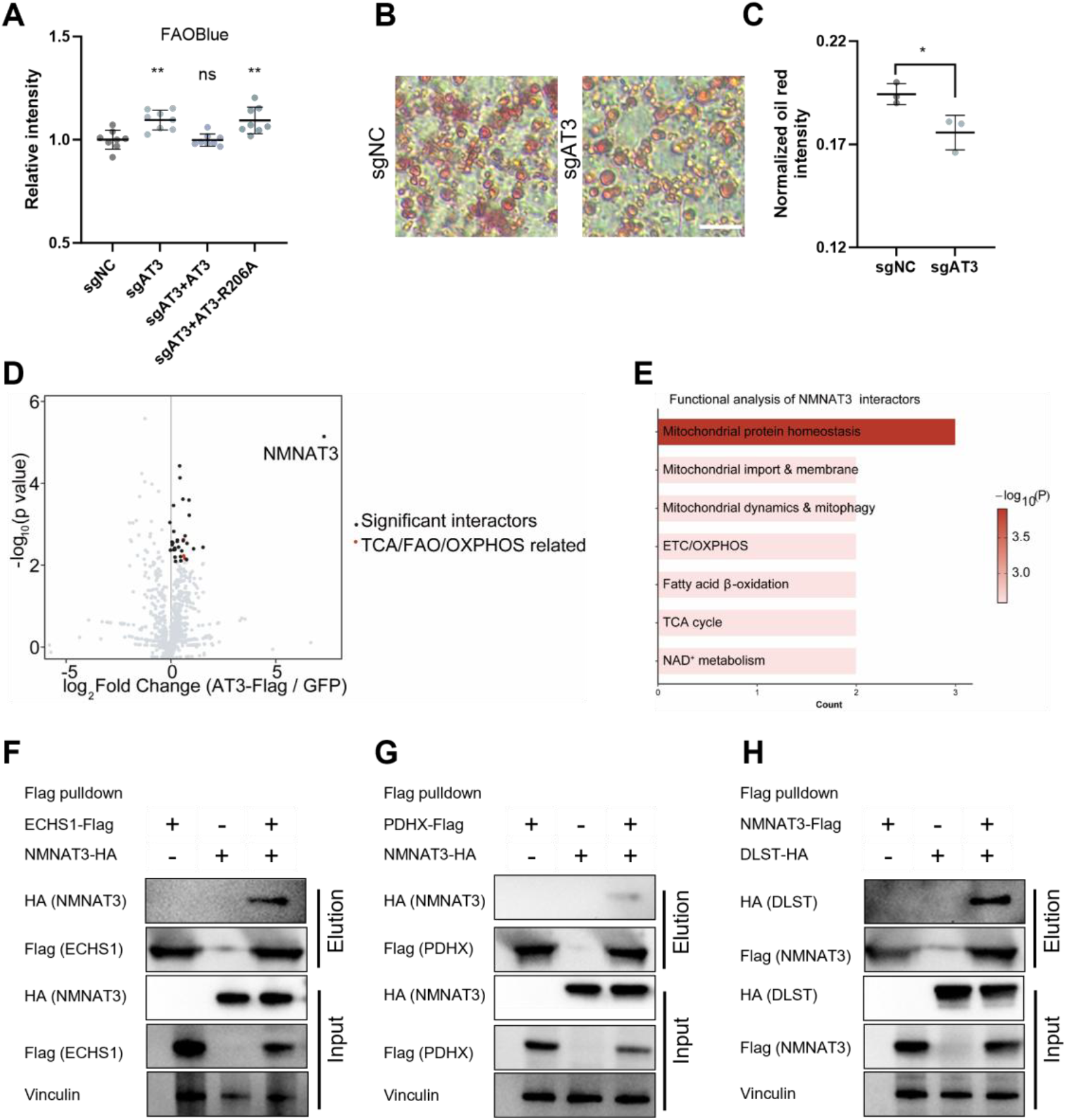
NMNAT3 modulates mitochondria metabolism and interacts with central carbon metabolic enzymes. (A) Quantitative analysis of mitochondrial FAO activity using the FAOBlue fluorescent probe in control and NMNAT3 knockdown HepG2 cells. (B) Representative microscopy images of Oil Red O staining showing neutral lipid accumulation in HepG2 cells. Scale bar, 15 μm. (C) Quantification of oil red o staining intensity corresponding to (B). (D) Volcano plot of affinity purification mass spectrometry results identifying potential NMNAT3 interacting proteins. Significantly enriched proteins (highlighted) include components of the TCA cycle, FAO, and OXPHOS complexes. (E) Functional analysis of NMNAT3 associated proteins, highlighting significant functional clusters related to mitochondrial protein homeostasis and central metabolic processes. (F) Co-IP of NMNAT3-HA with ECHS1-Flag in HEK 293T cells. (G) Co-IP of NMNAT3-HA with PDHX-Flag in HEK 293T cells. (H) Co-IP of NMNAT3-Flag with DLST-HA in HEK 293T cells. Data are presented as mean ± SD. Statistical significance was determined using one-way ANOVA analysis followed by Dunnett’s multiple comparisons test or two-tailed Student’s t-test. ns, P ≥ 0.05; *, P < 0.05, **, P < 0.01.

To uncover the underlying mechanism, we mapped the NMNAT3 interactome via affinity purification mass spectrometry (AP-MS). This revealed a specific network of mitochondrial proteins (Fig. 5D), with functional analysis highlighting central carbon metabolism pathways, including the TCA cycle, FAO, and OXPHOS (Fig. 5E). Co-immunoprecipitation (Co-IP) confirmed the association of NMNAT3 with key metabolic enzymes across distinct nodes: ECHS1 (FAO), PDHX (pyruvate to TCA transition), and DLST (TCA cycle) (Fig. 5F-H).

These interactions position NMNAT3 as a localized NAD^+^ buffering node adjacent to central metabolic machineries. Rather than driving net synthesis, NMNAT3 mediated hydrolysis acts analogously to SELO^13^. It spatiotemporally tunes the local NAD^+^/NMN ratio, thereby preventing metabolic gridlock and safeguarding mitochondrial homeostasis.

### HINT2 modulates the mitochondrial NAD^+^/NMN ratio

The mechanisms that set the mitochondrial NAD^+^/NMN ratio remain unclear. To identify potential regulators of mitochondrial NMN, we performed a targeted screen of literature-reported candidate genes using NMN biosensor. HINT2 and SLC25A48 both altered NMN levels (Fig. S10A), we focused on HINT2 here.

HINT2 knockdown in HepG2 cells lowered mitochondrial NMN and raised NAD^+^ (Fig. 6A-C). Overexpression had the opposite effect. HINT2 lacks intrinsic NAD^+^ hydrolase activity, so we asked whether it regulates NMNAT3. I*n vitro* assays with purified proteins showed that HINT2 markedly enhanced NMNAT3 hydrolysis of NAD^+^ (Fig. S6B, 5E, F). It did not affect NMNAT3 synthesis (Fig. 6D). A catalytically inactive HINT2 mutant (D80A) failed to stimulate hydrolysis *in vitro* and in cells (Fig. 6D-H). This interaction is therefore functionally specific. To test whether HINT2 act through NMNAT3 in cells, we knocked down NMNAT3. Neither HINT2 knockdown nor overexpression changed matrix NMN or NAD^+^ in this background (Fig. 6I, J). HINT2 thus requires NMNAT3 to regulate mitochondrial NAD^+^/NMN dynamics.

**Figure 6.**
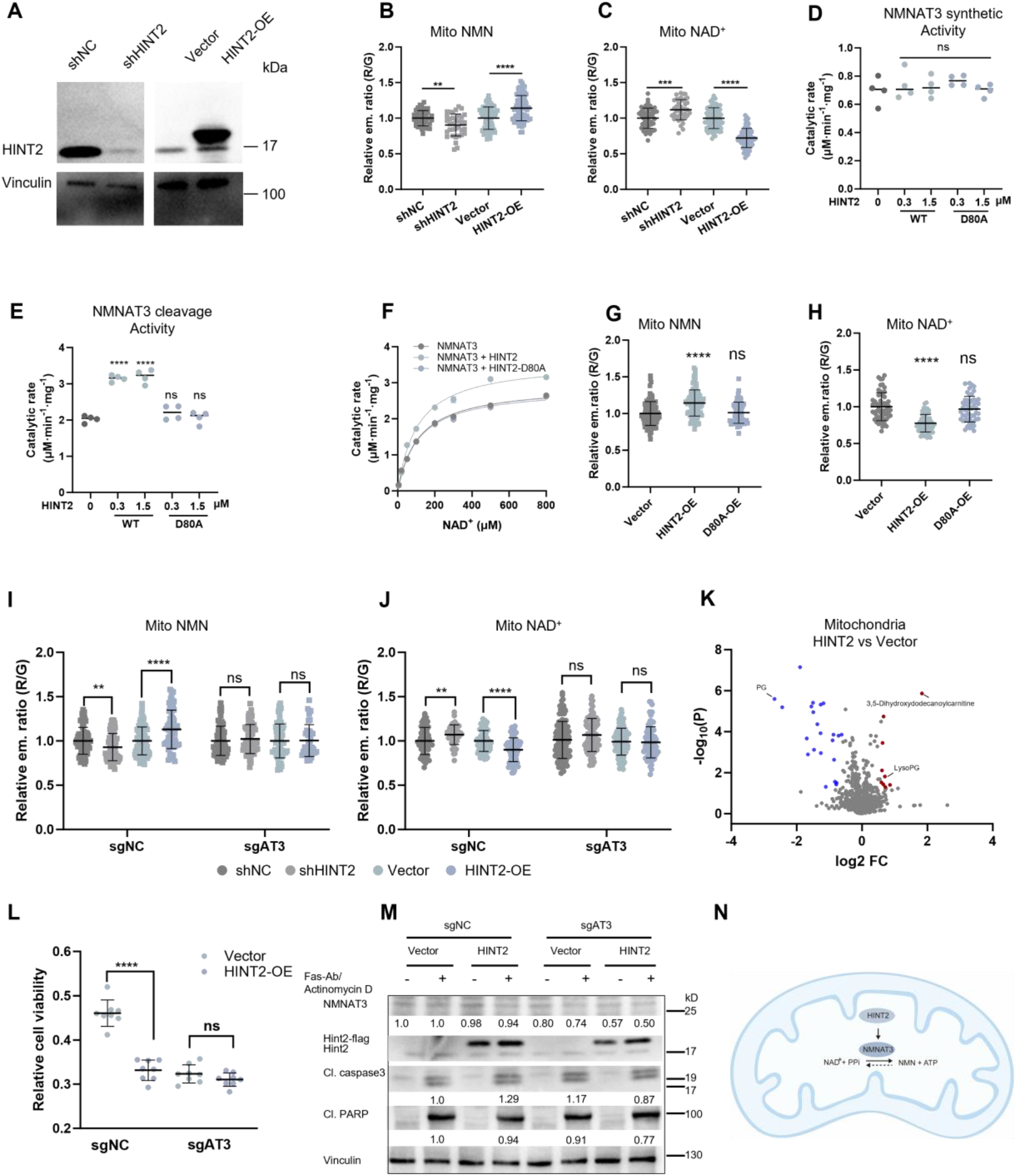
HINT2 enhances NMNAT3’s NAD^+^ hydrolyzing activity in mitochondria. (A) Immunoblotting analysis validating the knockdown and overexpression efficiency of HINT2 in HepG2 cells. (B and C) Relative emission ratios of mitochondrial NMN (B) and NAD^+^ (C) biosensors in HepG2 cells subjected to HINT2 knockdown or overexpression. (D and E) In vitro enzymatic analysis measuring the NAD^+^ synthesizing (D) and NAD^+^ hydrolyzing (E) catalytic rates of purified NMNAT3 in the absence or presence of HINT2. (F) Michaelis-Menten kinetics of NMNAT3 mediated NAD^+^ hydrolysis evaluated under varying NAD^+^ concentrations, supplemented with or without HINT2. (G and H) Overexpression of the catalytically inactive HINT2-D80A mutant did not alter mitochondrial NMN (G) or NAD^+^ (H) levels. (I and J) Evaluation of relative mitochondrial NMN (I) and NAD^+^ (J) levels in control or NMNAT3-knockdown HepG2 cells subjected to concurrent HINT2 knockdown or overexpression. (K) Volcano plot of untargeted metabolic analysis of purified HepG2 mitochondria under HINT2 overexpression or vector control. (L) Relative cell viability of HepG2 cells with control or NMNAT3-knockdown background, transduced with empty vector or HINT2. Cells were incubated 16 h with anti-Fas antibody (100 ng/mL) and actinomycin D (50 ng/mL) in. Viability was assessed by resazurin-based viability assay kit. (M) Immunoblotting analysis of apoptotic markers, NMNAT3, and HINT2 in control or NMNAT3 knockdown HepG2 cells. Cells were treated with or without Fas-antibody/Actinomycin D to induce apoptosis. (N) Proposed schematic model depicting how HINT2 acts as an upstream activator that specifically enhances NMNAT3 mediated NAD^+^ hydrolysis in mitochondria. Data are presented as mean ± SD; statistical significance was determined using one-way ANOVA analysis followed by Dunnett’s multiple comparisons test or two-tailed Student’s t-test. ns, P ≥ 0.05; *, P < 0.05; **, P < 0.01; ***, P < 0.001; ****, P < 0.0001.

To assess the metabolic consequences of HINT2 driven NMNAT3 activation, we performed untargeted metabolomics on immunopurified mitochondria. The volcano plot revealed HINT2 overexpression suppresses mitochondrial FAO, indicated by accumulation of 3,5-dihydroxydodecanoylcarnitine and decreased PG (Fig. 6K).

HINT2 suppresses tumor growth in several cancer lines, including HepG2^30–32^. We asked whether the growth inhibition relies on NMNAT3. HINT2 overexpression substantially restricted HepG2 cell viability. This suppression vanished when NMNAT3 was knocked down (Fig. 6L), even though NMNAT3 loss itself moderately reduced HepG2 viability. HINT2 overexpression also raised cleaved caspase-3, a marker of apoptosis. NMNAT3 knockdown reversed this increase (Fig. 6M).

Collectively, these findings establish a regulatory paradigm in which mitochondrial HINT2 enhances NMNAT3’s NAD^+^-hydrolytic activity. This interaction fine-tunes the mitochondrial NAD^+^/NMN balance and might contribute to the tumor-suppressive function of HINT2 (Fig. 6N).

### SLC25A48 regulates mitochondrial NMN homeostasis

Consistent with previous reports^14^, exogenous NAD^+^ restored matrix NAD^+^, but exogenous NMN failed to enter the matrix (Fig. S11A, B). SLC25A48 emerged from our screen as a regulator of mitochondrial NMN. It was reported to control mitochondrial choline import and metabolism^33,34^. SLC25A48 knockdown raised NMN in mitochondrial matrix and lowered NAD^+^ (Fig. 7A, B). Overexpression lowered matrix NMN and raised NAD^+^ (Fig. 7A, B).

**Figure 7.**
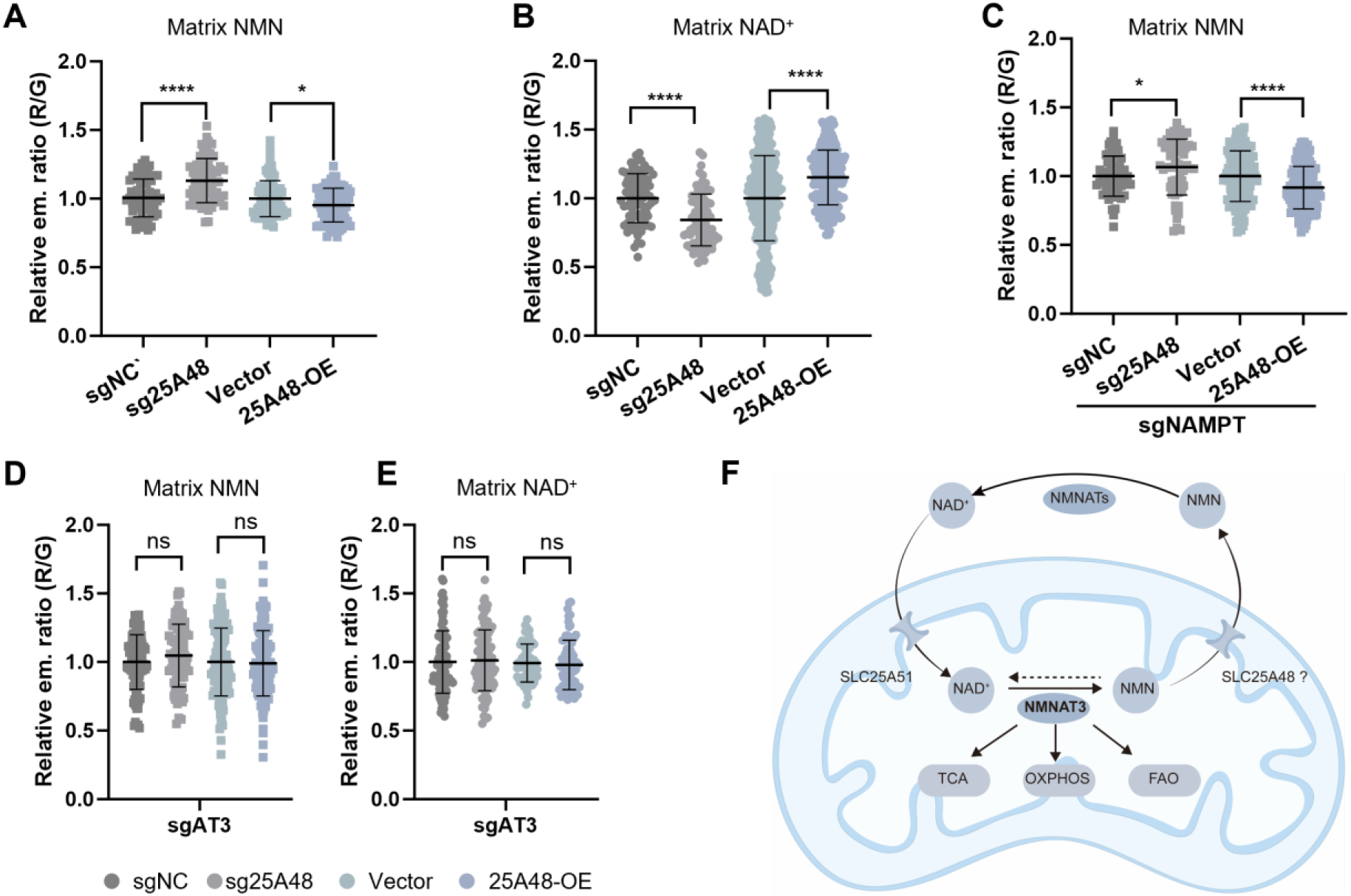
SLC25A48 regulates mitochondrial NAD^+^/NMN balance. (A) Relative emission ratios of compartment-specific biosensors measuring matrix NMN upon SLC25A48 knockdown or overexpression. (B) Relative emission ratios of compartment-specific biosensors measuring matrix NAD^+^ upon SLC25A48 knockdown or overexpression. (C) Relative emission ratios indicate matrix NMN levels in NAMPT knockdown cells subjected to SLC25A48 knockdown or overexpression. (D, E) Relative matrix NMN (D) and NAD^+^ (E) pools in NMNAT3 knockdown cells following SLC25A48 perturbation. (F) Proposed schematic model of the mitochondrial NAD^+^/NMN regulatory circuit: SLC25A51 imports NAD^+^, NMNAT3 hydrolyzes NAD^+^ to NMN, and SLC25A48 modulates the compartmental redistribution of the generated NMN to prevent metabolic gridlock. Data are presented as mean ± SD; statistical significance was determined using two-tailed Student’s t-test. ns, P ≥ 0.05; *, P < 0.05; ****, P < 0.0001.

Although mitochondrial NMN is most possibly generated locally via NAD^+^ hydrolysis, we wanted to exclude effects from cytosolic NMN import. We therefore knocked down NAMPT to block cytosolic NMN synthesis (Fig. S11C). Under this condition, SLC25A48 perturbation still changed matrix NMN in the same directions (Fig. 7C). SLC25A48 thus controls intrinsic mitochondrial NMN, not import. This regulation required endogenous NMN production: NMNAT3 knockdown abolished all effects of SLC25A48 on matrix NMN and NAD^+^ (Fig. 7D, E). Together, these results show that SLC25A48 regulates mitochondrial NMN levels.

## DISCUSSION

Here we report FrNADS and FrNMNS^1.0^, genetically encoded FRET sensors for NAD^+^ and NMN. FrNADS pairs StayGold-E138D with mScarlet-I3 and achieved a 3.6-fold emission ratio change for NAD^+35–37.^ FrNMNS^1.0^ achieves a 2.6-fold change for NMN. These values substantially exceed the typical dynamic range (< 1.5-fold) of prior FRET sensors^38^. Several genetically encoded NAD^+^ sensors have been developed previously, including LigA-cpVenus^21^, NAD-Snifit^22^, FiNad^23^, NS-X^24^ and ChemoX-NAD^+25^. LigA-cpVenus is pH-sensitive and cross-reacts with NMN and NR. FiNad offers an excellent dynamic range but remains sensitive to pH and NADH interference. NAD-Snifit and ChemoX-NAD^+^ require exogenous fluorophores, which complicates live cell work. FrNADS is insensitive to Ca^2+^ and Mg^2+^, interferences that prior sensors did not evaluate systematically. FrNADS also resists adenine nucleotides and remains stable across the physiological pH range (Fig. 1, S2). A summary table was presented in Table S1. We additionally developed FrNADS^ox^, a variant that resists oxidation, enabling NAD^+^ imaging in the ER and GA. FrNMNS^1.0^ complements our previously reported bioluminescent NMN sensor NMoRI^26^. Together, these tools extend our biosensor platform to live cell subcellular imaging and enable comprehensive analysis of NAD^+^ biology at organelle resolution.

Høyland et al. showed that subcellular NAD^+^ pools are interconnected and buffered by mitochondrial NAD^+8^. Steady-state bulk metabolomics cannot resolve the spatiotemporal dynamics of these connections. Previous work suggested that mitochondrial release NAD⁺ into the nucleus during oxidative stress to support PARP activity^27^. Our real-time imaging following MNNG treatment revealed nuclear NAD^+^ drops sharply and then recovers partially (Fig. 2B). This recovery was abolished completely by FK866, a NAMPT inhibitor (Fig. 2D). These data indicate that local salvage synthesis drives nuclear replenishment under acute genotoxic stress. We did not test whether blocking mitochondrial NAD⁺ import impairs this recovery; we therefore cannot exclude a parallel or backup contribution from mitochondria, particularly under prolonged or severe stress.

Subcellular compartmentalization allows cells to maintain distinct metabolic pools and balance consumption with biosynthesis^12^. SLC25A51 is established as the primary mitochondrial NAD^+^ transporter^15–17^. By contrast, the mechanisms that govern NAD^+^ homeostasis in peroxisomes, ER and GA remain largely unknown. We found that NUDT12 controls peroxisomal NAD^+^ and NMN levels directly (Fig. 3). This establishes NUDT12 as a key intrinsic regulator of the local NAD^+^/NMN balance. SLC25A17 knockdown markedly reduced peroxisomal NAD^+^ levels, while its overexpression elevated them (Fig. 3A-B). This provides direct functional evidence that SLC25A17 promotes NAD^+^ accumulation in peroxisomes, consistent with its proposed role as a peroxisomal NAD^+^ transporter^28^. These findings establish that peroxisomal NAD^+^ homeostasis reflects a balance between import mediated by SLC25A17 and consumption driven by NUDT12.

Mitochondria are central to energy metabolism, and mitochondrial NAD^+^ has been extensively studied as a redox and signaling cofactor. Both depletion and hyperaccumulation of the mitochondrial NAD^+^ pool are harmful^13^. This implies the existence of complex regulatory mechanisms. SLC25A51 is established as the primary transporter that sets the mitochondrial NAD^+^ pool size^15–17^. Goyal et al. showed that the mitochondrial NAD^+^ gradient is sustained by membrane potential and transport dynamics^39^. The physiological role of NMNAT3 remains controversial. Some studies found it dispensable *in vivo*^18^. Others reported benefits from overexpression^19,40,41^. Early biochemical hints suggested mitochondria localized isoform FKSG76 (NMNAT3) might cleave NAD^+^ rather than synthesize it^29^. Our sensors directly demonstrate that NMNAT3 mainly acts as a hydrolase in mitochondria (Fig. 4). NMNAT3 overexpression elevated mitochondrial NMN while depleting NAD^+^. Its knockdown produced the opposite effects. We establish NMNAT3 as an enzyme that hydrolyzes NAD^+^ in mitochondria and contributes to the local NMN pool. Jia et al. showed that SELO hydrolyzes NAD^+^ to NMN and AMP^13^. Its activity is pH responsive and coupled to fatty acid oxidation. Together, NMNAT3 and SELO form a redundant but differentially controlled layer of mitochondrial NAD^+^ buffering.

Crucially, recognizing NMNAT3 as a localized hydrolase reveals its true physiological function in regulating lipid metabolism. Contrary to the overexpression phenotype^19^, our genetic perturbations demonstrate that NMNAT3 knockdown promotes FAO activity (Fig. 5A-C). This indicates that endogenous NMNAT3 acts as a physiological brake on lipid catabolism. This concept is solidified by our AP-MS interactome, which reveals that NMNAT3 physically associates with central FAO and TCA enzymes, including ECHS1, PDHX, and DLST (Fig. 5D-H). Similar to SELO^13^, by tethering to these metabolic machineries, NMNAT3 creates localized zones of NAD^+^ depletion and NMN accumulation, thereby spatiotemporally restricting the activity of these NAD^+^ consuming engines.

Despite this local production, exogenous NMN does not elevate the NAD^+^ pool in isolated mitochondria^14,17^, suggesting NMN is not imported efficiently into the matrix. To identify additional regulators of mitochondrial NMN levels, we screened for candidates and identified HINT2 and SLC25A48. HINT2 has been linked to protein acetylation and mitochondrial NAD^+^ homeostasis^42^. Our results show that HINT2 lacks intrinsic NAD^+^ hydrolase activity but enhances the hydrolytic activity of NMNAT3 *in vitro* and in cells (Fig. 6D-H). Whether this enhancement reflects a direct physical interaction, allosteric activation, or an indirect effect remains to be determined. This regulatory interaction fine-tunes the mitochondrial NAD^+^/NMN balance and contributes to HINT2’s tumor-suppressive effects in HepG2 cells (Fig. 6K-M). HINT2 has also been reported to promote OXPHOS through interaction with MTCO2^43^. How an enhancer of NAD^+^ hydrolysis simultaneously supports oxidative phosphorylation remains unclear. We speculate that HINT2 engages distinct protein partnerships that vary with cellular context.

Finally, to avert localized metabolic gridlock from continuous NMN generation, SLC25A48 functions as a critical regulatory node for NMN clearance. SLC25A48 was recently identified as the mitochondrial choline transporter^33,34^. Our data show that SLC25A48 perturbation bidirectionally alters matrix NMN levels, suggesting a functional link between choline import and NMN homeostasis. Definitive elucidation of SLC25A48’s molecular mode of action will require liposome reconstitution or high-resolution structural analysis.

Together, our findings delineate a regulatory network in which NMNAT3 generates NMN from NAD^+^, HINT2 augments its hydrolytic activity, and SLC25A48 modulates NMN compartmental redistribution (Fig. 7F). This network prevents excessive NMN accumulation and maintains NAD^+^ homeostasis. Fitzpatrick and Kory described mitochondria as a rheostat for cellular NAD^+^ levels^12^. Our data extend this model by revealing how local hydrolysis and efflux cooperate to fine-tune the mitochondrial NAD^+^ pool. This intricate balance has implications for NAD^+^-boosting therapies: supplementation with NAD^+^ precursors may be counterproductive in tissues with high mitochondrial NMNAT3 activity, as it could promote matrix NMN accumulation rather than elevate NAD^+^, especially when NMN clearance is impaired.

The mechanisms that regulate the mitochondrial NAD^+^ pool are becoming clearer and involve SLC25A51, NMNAT3, SELO, HINT2, and SLC25A48, as well as additional yet-to-be-identified regulators. By contrast, how cells adjust NAD^+^ levels in peroxisomes, the ER, and the GA remains largely unknown. Our biosensor toolkit extends detection capability to previously inaccessible organelles. We anticipate that FrNADS and FrNMNS^1.0^ will facilitate studies of compartmentalized NAD^+^ metabolism in health and disease.

### Limitations of the study

First, biosensor measurements report relative, not absolute, NAD^+^ and NMN concentrations. Second, the conclusion that nuclear NAD^+^ replenishment relies on local salvage rests on pharmacological inhibition. Third, although HINT2 modulates NMNAT3 activity, a direct physical interaction has not been demonstrated, and allosteric or indirect mechanisms remain possible. Fourth, dedicated reconstitution and structural studies are required to determine the mechanism of SLC25A48. Finally, our findings are derived from immortalized cell lines and await validation in primary cells and *in vivo* models.

## Methods

### Molecular biology

Primers for cloning were obtained from TSINKE Biotech, and PCRs were performed using the KOD one master mix (Toyobo) according to the manufacturer’s protocol. Sensor variants were generated by site-directed mutagenesis with specific oligonucleotide primers purchased from TSINKE Biotech. Seamless cloning was used as the standard method for cloning. Transformations were performed using standard heat shock transformation. Site-directed mutagenesis was performed using overlap PCR. All plasmids were amplified in the *Escherichia coli* strain stbl3. DNA sequences were validated by Sanger sequencing (TSINKE Biotech), and DNA was stored at −20 ℃ until further use.

A pET51b vector was used as the backbone for protein expression in *E. coli* BL21 (DE3). Target genes were flanked by an N-terminal Strep-tag II and C-terminal His tag for protein purification. A pCDH-CMV-MCS-EF1-Neo vector or pLV3-CMV-MCS-Puro/Blast vector was used for gene expression in mammalian cells. For targeted expression at subcellular locations, the targeted gene was flanked by N- and/or C-terminal localization sequences. Localization sequences:

- Cytosol, nuclear export signal (NES) (ATGCTGCAGAATGAACTGGCACTGAAGCTCGCAGGCCTGGACATTAACAAGACCGGTGGAAGCGGC),
- Nucleus, nuclear targeted sequence (NTS) (ATGGATCCAAAAAAGAAGAGAAA GGTA),
- Mitochondrial matrix, 2 x COXVIII (ATGTCTGTTCTGACTCCTCTGCTGCTCCGGGGTCTCACAGGTTCCGCAAGAAGACTCCCCGTGCCTAGGGCCAAAATTCATTCACTGGGGGACCCCATGAGCGTGCTCACCCCACTCCTGCTGCGGGGGCTGACCGGCAGCGCTAGGCGGCTGCCAGTCCCCAGGGCCAAGATCCACA GTCTCGGCGATCCCAAG)
- Peroxisome, peroxisomal targeting signal 1 (PTS1) (LGRGRRSKL),
- Endoplasmic reticulum, signal sequence of calreticulin (MLLSVPLLLGLLGLAVAD RS) in N-terminal and KDEL in C-terminal,
- Golgi apparatus, signal sequence from MGAT2 (MRFRIYKRKVLILTLVVAACGFV LWSSNGRQRKNEALAPPLLDAEPARGAGGRGGDHPSVAVGIRRVSNVSAASLVPAVPQPEADNLTLLDPGGSGGS).

### Protein expression and purification

*E. coli* BL21 (DE3) carrying the expression plasmid was grown in 200 mL LB medium containing 50 μg/mL ampicillin at 37 ^◦^C until the cultures reached an OD_600_ of approximately 0.6. The growth temperature was then shifted to 18 ^◦^C and protein expression was induced by adding 0.2 mM isopropyl-β-D-thioacetamide (IPTG) for 16 h. Bacteria were harvested by centrifugation at 4,000 × g for 10 minutes at 4 ^◦^C and suspended in 50 mM Tris-HCl buffer (pH 8.0) containing 500 mM NaCl and 10 mM imidazole. Cells were lysed via high pressure homogenizer. Cell lysate was centrifuged at 12,000 × g for 10 min at 4 ^◦^C, and the supernatant was loaded sequentially onto Ni-NTA (Smart Lifesciences Inc., China) and Strep-Tactin columns (Smart Lifesciences Inc., China) for purification. The purified protein was desalted and exchanged into 50 mM HEPES buffer containing 50 mM NaCl (pH 7.2) with an Amicon Ultra-15 centrifugal filter (10 kDa MWCO, Merck Millipore Inc.). Protein concentration was determined by BCA assay and subsequently diluted to the requisite concentration for subsequent *in vitro* assays. Protein purity was confirmed by standard SDS-PAGE. Purified proteins were stored in 50% (v/v) glycerol at −20 ^◦^C until further use.

### Sensor titration

Fluorescence measurements were performed in 100 μL of HBS buffer (20 mM HEPES, 150 mM NaCl, pH 7.2) or HPS buffer (20 mM HEPES, 150 mM NaCl, 2 mM MgCl_2_, 2 mM CaCl_2_, 2.5 mM KCl, pH 7.2) supplemented with 0.1 mg/mL BSA in black non-binding, flat-bottom 96-well plates (SPL). Excitation wavelength was set to 488 nm, emission was measured at 520 nm and 595 nm using a FlexStation 3 Multi-Mode microplate reader. The emission intensity ratios between 595 nm and 520 nm were plotted against compound concentrations. The sensor’s maximum ratio (R_max_), minimum ratio (R_min_), c50 and Hill coefficient (h) were obtained by fitting the data to the Hill-Langmuir equation:

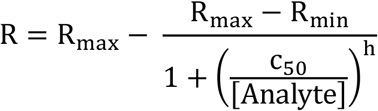

For substrate titration, the sensor protein was diluted to 200 nM in HBS or HPS buffer. Analyte solutions were prepared at 10x final concentration in water and diluted to 1x in the assay wells. Sensor titration in the presence of structurally similar analytes was performed as described above but in the presence of the competing analyte.

The pH sensitivity of the sensors was evaluated using two buffers (SPG pH 4.0 and 10.0, both 1M; Jena Bioscience) mixed in defined ratios to yield buffers with pH values ranging from 6.8 to 8.0. The buffers with varied pH were used to replace HEPES in the assay system.

### Cell culture

HEK 293T (ATCC, CRL-3216) and HepG2 (Immocell, IM-H038) cells were cultured in DMEM (11885, Gibco) supplemented with 10% FBS (VivaCell C04001) at 37 ^◦^C with 5% CO_2_ in a humidified cell culture incubator. Cells were handled under a sterile laminar flow hood and kept in culture for a maximum 15 passages. Contamination with *mycoplasma* was regularly checked by PCR.

### Generation of NAD^+^ and NMN sensor cell lines

Lentiviral constructs (LentiCRISPR, PCDH and pLV3) were transfected into 293T cells together with pMD2G and psPAX2 for lentiviral production. All transfections were done by liposome-based transfection. At 48 h post transfection, the medium was collected and filtered through a 0.22 μm PES membrane. The resulting viral solution was then divided into aliquots, flash frozen, and stored at −80 ^◦^C until further use.

For cell infection, HEK 293T and HepG2 cells cultured in 60 mm plates were exposed to the virus solutions at an MOI approximately ten. Culture media were replaced 6 h after infection. After 48 h incubation period post-infection, cells were harvested and subjected to serial dilution to select for infected cells, using sensors fluorescence as a selection marker.

### Construction of gene knockout, knockdown and overexpression cell lines

CRISPR-Cas9-mediated ablation of *NMNAT3* gene in HEK 293T cells was achieved with CRISPR-Cas9 RNPs with the help of Ubigene Biosciences (Guangzhou, China). The Cas9 protein (Ubigene, YK-Cas9-50) and gRNA were incubated at room temperature for 10-20 min, followed by electroporation via a Neo transfection system according to the manufacturer’s instructions. After 2 days, the cells were sorted into 96-well plates via cell sorter to form single clones. Candidate clones were screened via genomic sequencing to verify the presence of mutations in both alleles.

To create shRNA for target gene knockdown, they were ligated into a pLKO.1 puro vector and produce lentivirus as described. Sensor expressing cells were transfected with this lentivirus to induce the knockdown of targeted genes. The specific shRNA sequences used for knockdown are as follows: shPXMP2, GCCTCTGAGATATGCCGTTTA; shNUDT12, GCTTAGCTCTAGCAGTGTCTA; sh25A17 (SLC25A17), GCTCTCATGTTCCTTGTTTAT; shHINT2, CCGACTTGTGATCAACGATGG. NMNAT3, NAMPT and SLC25A48 were knocked down by using the following sgRNA target sequences: sgNMNAT3, CTGGAAGGATGCGCACATCC; sgNAMPT, CCGGCCCGAGATGAATCCTG. Non-targeting control sgNC sequence: GGTTCTCCGAACGTGTCACGT; sg25A48: CCTGGACTTCAAACCGAGGT. The sgRNA were ligated into LentiCRISPR v2 vector. HEK 293T cells were transfected with this plasmid to generate lentivirus. Subsequently, cells were transfected with this lentivirus to induce the knockdown of targeted gene. Lentivirus derived from shRNA, sgRNA, or overexpression plasmids, supplemented with 8 μg/mL polybrene, were added to the respective cells for 24 hours, followed by medium exchange, recovery and selection or downstream experiments. Cells were selected either by antibiotic treatment (puromycin, 2-6 μg/mL; blasticidin, 10 μg/mL). Transduced cell pools were validated by western blot.

### Western blot and immunoprecipitation

Cells were washed three times with ice-cold PBS and lysed in RIPA buffer with protease and phosphatase inhibitors. Protein samples were electrophoresed on 4-20% precast gel and transferred to a 0.45 μm PVDF membrane. Membranes were blocked with 5% non-fat milk in Tris-buffered saline containing 0.1% (v/v) Tween-20 (TBST). Primary antibodies were diluted in 1% non-fat milk in TBST and incubated at 4 ^◦^C overnight. After three washes, the secondary antibody, diluted in TBST, was incubated for 1 h at RT. After another three washes, target proteins were visualized with iBright imaging system. The following primary and secondary antibodies were used for WB analysis: PXMP2 (Proteintech, 24801-1-AP, 1:1000), NUDT12 (Proteintech, 17487-1-AP, 1:1000), SLC25A17 (ABclonal, A14840, 1:1000), HINT2 (Proteintech, 17986-1-AP, 1:1000), Caspase 3 (Proteintech, 19677-1-AP, 1:1000), PARP (Cell Signaling Technology, 9542, 1:1000), NAMPT (Proteintech, 11776-1-AP, 1:1000), Vinculin (Cell Signaling Technology, 13901, 1:2000), NMNAT3 (Santa Cruz, sc-3904 33, 1:500), SLC25A51 (ABclonal, A21105, 1:1000), anti-Flag M2 antibody (Millipore, F1804), HA tag (Proteintech, 51064-2-AP), anti-Rabbit IgG (Biosharp, BL003A, 1:10000), anti-Mouse IgG (Biosharp, BL021A, 1:10000).

For immunoprecipitation (IP) with transfected HEK 293T cells, cells were lysed in IP lysis buffer (PBS, 0.5% Triton X-100, phosphatase inhibitor and proteinase inhibitor cocktail) for 1 h. The suspensions were centrifuged at 12,000 × g for 10 min at 4 ^◦^C. Part of the supernatant (20 μL) was transferred to a new tube and added SDS-PAGE loading buffer to make input sample. The remaining supernatant was incubated with 10 μL Flag M2 magnetic beads (Merck, M8823) overnight with gentle rotation at 4 ^◦^C. Subsequently, the beads were washed with IP wash buffer (PBS, 0.1% Triton X-100, phosphatase inhibitor and proteinase inhibitor cocktail) extensively. The bound proteins were then eluted with PBS containing 0.5 mg/mL 3 x Flag peptide. The eluted proteins were then subjected to western blot analysis.

### Widefield microscopy

Widefield microscopy experiments were performed on an Olympus IX83 microscope equipped with Uplan Apo 40x/0.95-NA and 100x/1.50-NA oil objectives and an external filter wheel. Cells were washed three times with PBS, incubated in live cell imaging solution (Invitrogen) containing 10 mM glucose, and equilibrated on the microscope stage for 30 min. Images were captured as 16-bit with 250 ms exposure. For live-cell widefield FRET imaging, an LED was used for donor excitation via a GFP filter set. The emitted light was split by a 488/561 nm dichroic mirror into two channels: the GFP channel was captured by a CMOS camera, and the RFP channel was simultaneously captured by another CMOS camera.

### Image analysis

Image analysis was performed with FIJI^44^. Background signal was subtracted with a rolling ball radius of 50 pixels. The processed images were further used for quantitative analysis and figure generation. For the generation of FRET emission ratio images, the RFP channel was thresholding (Otsu method) to create mask and then applied to both GFP and RFP channels that were subsequently divided (RFP channel/GFP channel). Ratiometric images were represented with ‘fire’ look-up table. ROIs were manually defined for each cell, cells displaying unhealthy phenotypes were excluded from analyses.

### Fluorescence measurement of FAO activity

The cells were seeded in black 96-well plates, washed twice with Hanks’ balanced solution (HBS) and incubated with 20 μM FAOBlue (Funakoshi, FDV-0033T) in HBS for 30 min at 37 ℃. To evaluate background signal, FAOBlue was also added wells without cells. Fluorescence was measured with a FlexStation 3 Multi-Mode microplate reader (Ex/Em = 405/460 nm). To normalize measurements, protein concentration was measured with BCA.

### Oil Red O staining for analysis of lipid content

Oil Red O staining was used to assess the levels of neutral lipids accumulated in cultured cells. Cultured cells were grown in 6 well plate, and fixed in 4% paraformaldehyde for 20 min at room temperature, followed by a brief wash with water. Subsequently, cells were incubated with 60% isopropanol for 5 min before staining with oil red working solution at room temperature for 20 min. After Oil Red O staining, cells were washed with water and imaged with inverted microscope.

### Affinity enrichment mass spectrometry

HEK 293T cells expressing NMNAT3-Flag or GFP-Flag, were lysed in 500 μL cold IP lysis buffer for 1 h. Insoluble materials were pelleted and the supernatant was retained. The protein concentration was quantified by BCA, and equal masses of cell supernatant were mixed with 30 μL Flag M2 magnetic beads for 3 h at 4 ℃ with gentle rotation. Following incubation, beads were washed four times in IP wash buffer and twice in final wash buffer (20 mM HEPES, pH 7.4, 100 mM NaCl) before being subjected to on-bead trypsin digestion. The digested supernatant was dried under vacuum. 40 μL of 0.1% FA/H_2_O (formic acid in water) was added to the centrifuge tube containing the drained peptides, vortexing vigorously for 5 min to fully dissolve the peptides, centrifugation at 20,000 x g at 4 ℃ for 30 min, and 35 μL of the supernatant peptide solution was taken into a new centrifuge tube, waiting for the machine.

The peptides were re-dissolved in solvent A (A: 0.1% FA/H_2_O) and analyzed by Vanquish Neo UHPLC system (Thermo Fisher Scientific, MA, USA) coupled to Orbitrap Astral mass spectrometer (Thermo Fisher Scientific, MA, USA) with FAIMS. 200 ng peptide sample was loaded onto AUR3-15075C18-TS column (15 cm x 75 μm ID, 1.7 μm C18, Ionopticks Aurora Elite) and separated with 8 min-gradient starting at 8% buffer B (80% acetonitrile with 0.1% FA), followed by a stepwise increase to 17% in 1.5 min with the column flow rate of 1000 nL/min, 40% in 5 min with the column flow rate of 400 nL/min, 99% in 0.5 min with the column flow rate of 700 nL/min and stayed there for 1 min with the column flow rate of 1000 nL/min. The column temperature of 50 ℃. The mass spectrometer was run under data independent acquisition mode (DIA). For Orbitrap experiments, a survey scan was acquired at 380-980 m/z, the resolution was 240000, Normalized AGC target of 500% and a maximum injection time of 3ms, FAIMS CV was −42. For Astral experiments, the Precursor Mass range was 380-980 m/z, Normalized AGC target of 500% and a maximum injection time of 3 ms. The HCD collision energy was 25%. The isolation Window was 2 m/z and the overlap between every window was 0. Loop time was 0.6s. FAIMS CV was −42.

### NMNAT3 activity assay

The purified human recombinant NMNAT3 (UniProt Q96T66-1) was incubated with the indicated substrates in reaction buffer (50 mM Tris HCl, pH 8, 5 mM MgCl2, 200 mM NaCl). The reaction was run for 30 minutes at room temperature. The reaction was stopped by diluting the sample 1:100 with ice-cold water, and then the concentration of NAD^+^ was quantified by NAD^+^ sensor as described previously^24^. NAD^+^ [500 μM] and PPi [5 mM] were used as substrate for the cleavage activity assay, NMN [500 μM] and ATP [2 mM] were used as substrate for the synthetic activity assay. Final concentration of NMNAT3 was around 20 nM.

### Cell proliferation assay

For assessment of cell activity, 10,000 cells per well were seeded in the cavities of black 96-well plates. The following day, cells were washed with PBS and treated with corresponding treatments for another 24 h. Then, the cell viability was quantified using CellTiter-Blue kit (Promega) following the manufacturer’s protocol.

### Untargeted metabolic profiling of immunopurified mitochondria

Mitochondria were immunopurified from HepG2 cells expressing 3xHA-OMP25-GFP was conducted according to the protocol by Chen et al^45^. In brief, a total of 2 x 10^7^ cells were washed once in PBS, collected in cold PBS and centrifuged at 300 g for 10 min at 4 ℃. The cell pellet was resuspended in 2 mL KPBS, followed by homogenization with 30 strokes in a homogenizer containing a pure PTFE head (VWR international). Cell debris was spun down at 800 g for 10 min and supernatant was incubated with 200 μL anti-HA magnetic beads on a rotator shaker for 5 min at 4 ℃, followed by three washes with ice-cold KPBS. 5 μL of beads were lysed with RIPA buffer to quantification protein concentration for normalization. The remaining beads were extracted in 80% ice-cold methanol solution, with sonication and vortex. Incubated for 30 min at −20 ℃ to precipitate proteins, and centrifuged at 20,000 g for 10 min at 4 ℃, supernatant was centrifuged for 5 min again. The supernatant was transferred to a fresh vial for UPLC-HRMS analysis. The quality control sample was prepared by mixing an equal aliquot of the supernatant of samples. An ACQUITY UPLC HSS T3 column (100 mm x 2.1 mm, 1.8 μm, Waters) was used for separation. The mobile phase consists of phase A (5 mM/L ammonium acetate + 5 mM/L acetic acid + water) and phase B (acetonitrile). Gradient elution conditions were set as follows: 0-0.8 min, 2% B; 0.8-2.8 min, 2%-70% B; 2.8-5.6 min, 70%-90%B; 5.6-6.4 min, 90%-100% B; 6.4-8.0 min, 100% B; 8.0-8.1 min, 100%-2% B; 8.1-10.0 min, 2% B. The flow rate was 0.35 mL/min. The injection volume for each sample was 4 μL. The column oven was maintained at 40 ℃. A high-resolution tandem mass spectrometer triple TOF 6600 (AB SCIEX) was used to detect metabolites eluted from the column. Each sample was operated in both positive and negative electrospray ionization mode. ESI temperature was 500 ℃. The voltage is +5000 volts in positive ion mode and −4500 volts in negative ion mode. The curtain gas pressure of the ion source is 30 psi, Gas 1 (Auxiliary gas) and Gas 2 (Sheath gas) pressures were both set to 60 psi. The mass spectrometric data were obtained with full scan and information dependent acquisition (IDA) modes. In one acquisition cycle, the full scan acquisition range is 60-1200 Da, and the full scan acquisition time is 150 ms. Then, the top 12 signal ions with a signal accumulation intensity of more than 100 were selected from the full scan for IDA scanning, the IDA acquisition range is 25-1200 Da, and the acquisition time is 30 ms. Dynamic exclusion is set to 4 s.

### Mitochondrial isolation for imaging

Mitochondria were isolated as described previously^17^. Cells were cultured on a 100-mm plate and collected by trypsinization. Mitochondria were isolated by homogenizing cell in 2 mL of mitochondrial isolation buffer (210 mM mannitol, 70 mM sucrose, 10 mM HEPES, 1 mM EGTA, 0.25% fatty acid free BSA, pH adjusted to 7.2 with KOH) with 20 strokes in a dounce homogenizer containing a pure PTFE head (VWR international). Mitochondria were collected by differential centrifugation. Cell debris was spun down at 800 g for 10 min and supernatant was transferred to a new tube. This process was repeated until no cell debris pellet was present. Then the supernatant was spun at 11,000 g for 15 min. The mitochondria pellet was resuspended in 200 μL mitochondria incubation media [125 mM KCl, 10 mM MOPS (pH 7.2), 2 mM MgCl_2_, 2 mM KH_2_PO_4_, 10 mM NaCl, and 1 mM EGTA].

To image isolated mitochondria, the resuspended mitochondria were added into 8-chamber glass bottom slide (Cellvis). The suspension was added indicated compounds and sedimentated for 30 min at room temperature and then evenly covered with 0.2% low melting point agarose for widefield FRET imaging.

## Quantification and statistical analysis

All experimental results are expressed as Mean ± SD. Data comparisons between two groups were performed using unpaired two-tailed Student’s t-test. Comparisons among multiple groups were performed using one-way or two-way ANOVA followed by Dunnett’s multiple comparisons test, as specified in figure legends. A P value smaller than 0.05 was considered statistically significant.

## Acknowledgements

This work is supported by National Natural Science Foundation of China (22307135, 22577149), Research Projects on Key Areas of General Colleges and Universities, Department of Education of Guangdong Province (2022ZDZX2071), Shenzhen Science and Technology Program (JCYJ20220818100804009, ZDSYS20210623091810032), Natural Science Foundation of Guangdong Province (2023A1515010715), Shenzhen Institute of Advanced Technology, Chinese Academy of Sciences.

## Author Contributions

Conceptualization, Q.Y., and L.C.; Methodology, L.C.; Investigation, L.C., X.C., and Y.W.; Formal Analysis, L.C.; Writing-Original Draft, L.C.; Writing-Review & Editing, Q.Y., L.C.; Funding Acquisition, Q.Y., L.C.; Supervision, Q.Y.; Project Administration, Q.Y., L.C.

## Data and Code Availability

Plasmids and cell lines are available from the lead contact with a completed Materials Transfer Agreement.

