## Supplemental file for "Spatiotemporal biosensor profiling reveals an autonomous mitochondrial NAD^+^/NMN regulatory network centered on NMNAT3"

[3]Lead contact

**Table S1. Comparison of known genetically encoded $NAD^+$ sensors.**

| Sensor name | Type | Analogues decrease affinity | Physiological pH sensitivity | Affinity | Dynamic Ranges | Divalent ion interference | ATP/ADP interference | Reference |
|---|---|---|---|---|---|---|---|---|
| $NAD^+$-Snifit | BRET | No | Insensitive | 0.27 – 423 μM | ~ 7 | ND | Low | [1,2] |
| LigA-cpVenus | Intensiometric | Yes | Sensitive | ~ 65 μM | ~ 2 | ND | High | [3] |
| FiNad | Intensiometric | Yes | Sensitive | ~ 1300 μM | ~ 8 | ND | High | [4] |
| NS series | BRET/FRET | No | Insensitive | 47.4 nM - 3.1 mM | 1.8 - 3.1 | ND | High | [5] |
| ChemoX-NAD | BRET/FRET | Yes | Insensitive | 30 – 200 μM | 3 - 35 | ND | High | [6] |
| FrNADS | FRET | No | Insensitive | 30.2 μM | 3.6 | Low | Low | This study |

ND means not determined.

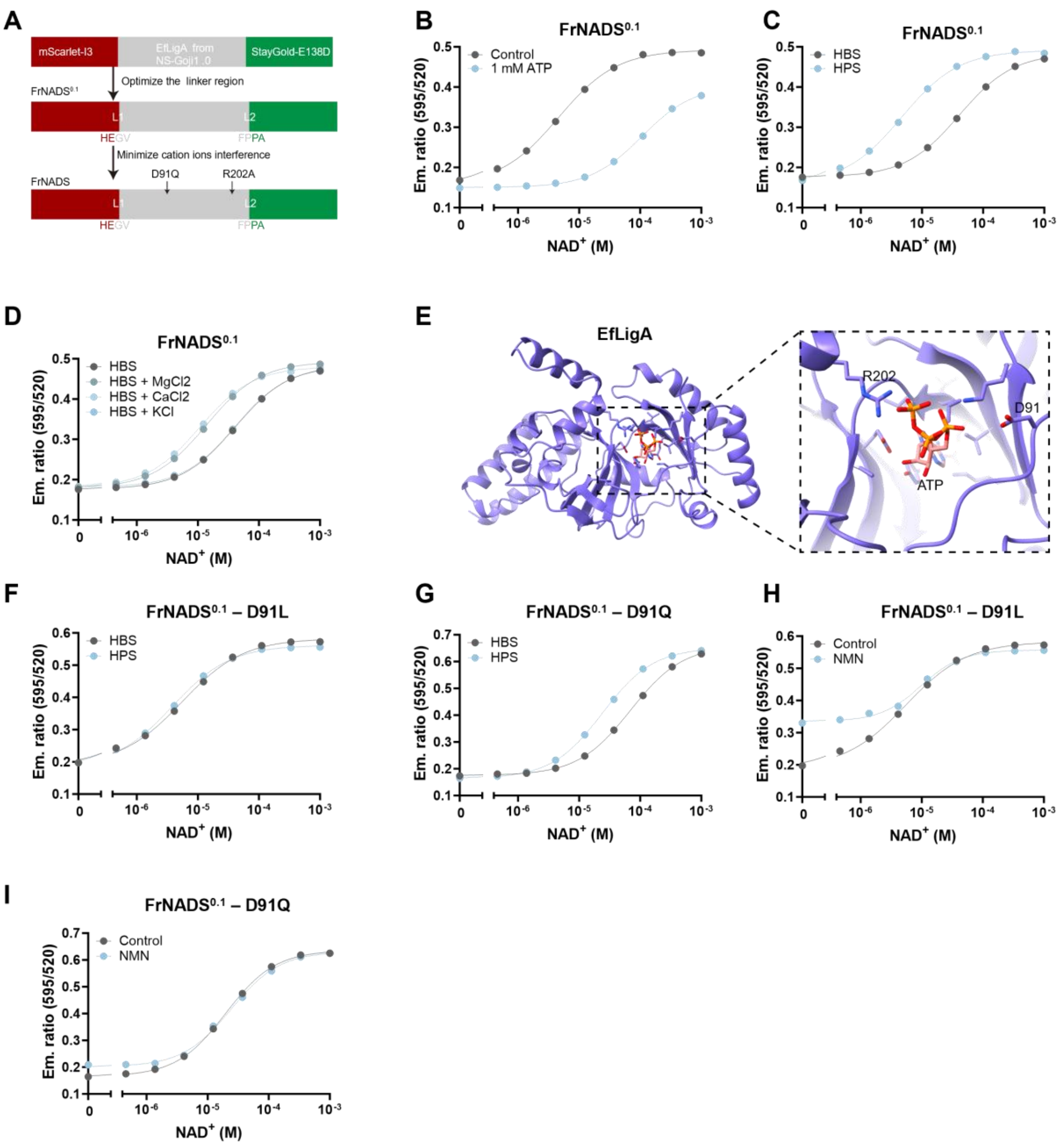


**Figure S1. Development of the NAD$^+$ sensor FrNADS, related to Figure 1.** (A) Schematic workflow of the development process for FrNADS. (B) Titration curves for FrNADS$^{0.1}$ with or without the presence of 1 mM ATP. (C) Titration curves of FrNADS$^{0.1}$ in HBS buffer (20 mM HEPES, 150 mM NaCl, pH 7.2) versus HPS buffer (20 mM HEPES, 150 mM NaCl, 2 mM $MgCl_2$, 2 mM $CaCl_2$, 2.5 mM KCl, pH 7.2), demonstrating cation-induced affinity shift. (D) Response curves of FrNADS$^{0.1}$ in HBS buffer supplemented with additional $MgCl_2$, $CaCl_2$, or KCl. (E) Structural model of ATP docked in the *Ef*LigA binding pocket (PDB code 1tae), highlighting D91 and R202 near the binding site. (F) Titration curves of the D91L variants in HBS versus HPS buffer. (G) Titration curves of the D91Q variant in HBS versus HPS buffer. (H) Titration curves of D91L variant with or without the presence of 50 μM NMN. (I) Titration curves of the D91Q variant with or without the presence of 50 μM NMN. Data are presented as mean ± SD from at least three independent measurements.

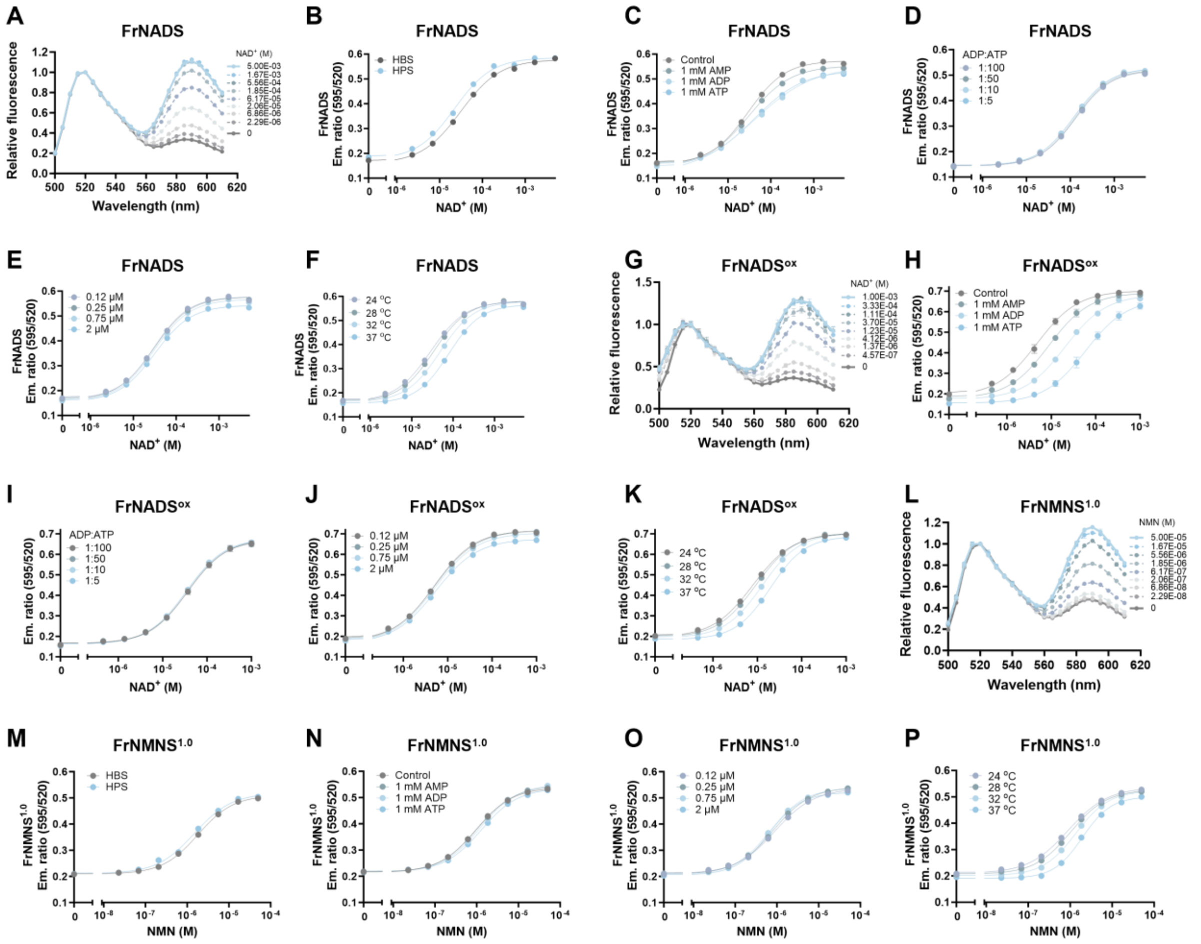


**Figure S2. Additional characterization of FrNADS, FrNADS$^{ox}$, and FrNMNS$^{1.0}$, related to Figure 1.** (A) Normalized emission spectra of FrNADS in response to increasing NAD$^+$. (B) Titration curves of FrNADS in HBS versus HPS buffer, confirming the absence of cation interference after optimization. (C) Titration curves of FrNADS to NAD$^+$ in the presence of 1 mM AXPs (AMP, ADP, ATP), showing minor interference. (D) Titration curves of FrNADS to NAD$^+$ under variable ADP:ATP ratios, indicating no effect on sensor performance. (E) Titration curves of FrNADS to NAD$^+$ at different sensor concentrations, demonstrating concentration independence. (F) Effect of temperature on the emission ratio of FrNADS, showing modest sensitivity. (G) Normalized emission spectra of FrNADS$^{ox}$ in response to increasing NAD$^+$ concentrations. (H) Titration curves of FrNADS$^{ox}$ to NAD$^+$ in the presence of 1 mM

AXPs. (I) Titration curves of FrNADS$^{ox}$ to NAD$^+$ under variable ADP/ATP ratios, indicating no effect on sensor performance. (J) Titration curves of FrNADS$^{ox}$ to NAD$^+$ at different sensor concentrations. (K) Effect of temperature on the emission ratio of FrNADS$^{ox}$. (L) Normalized emission spectra of FrNMNS$^{1.0}$ in response to increasing NMN concentrations. (M) Titration curves of FrNMNS$^{1.0}$ in HBS versus HPS buffer, confirming the absence of cation interference. (N) Titration curves of FrNMNS$^{1.0}$ to NMN in the presence of 1 mM AXPs. (O) Titration curves of FrNMNS$^{1.0}$ to NMN at different sensor concentrations. (P) Effect of temperature on the performance of FrNMNS$^{1.0}$. Data are presented as mean ± SD from at least three independent measurements.

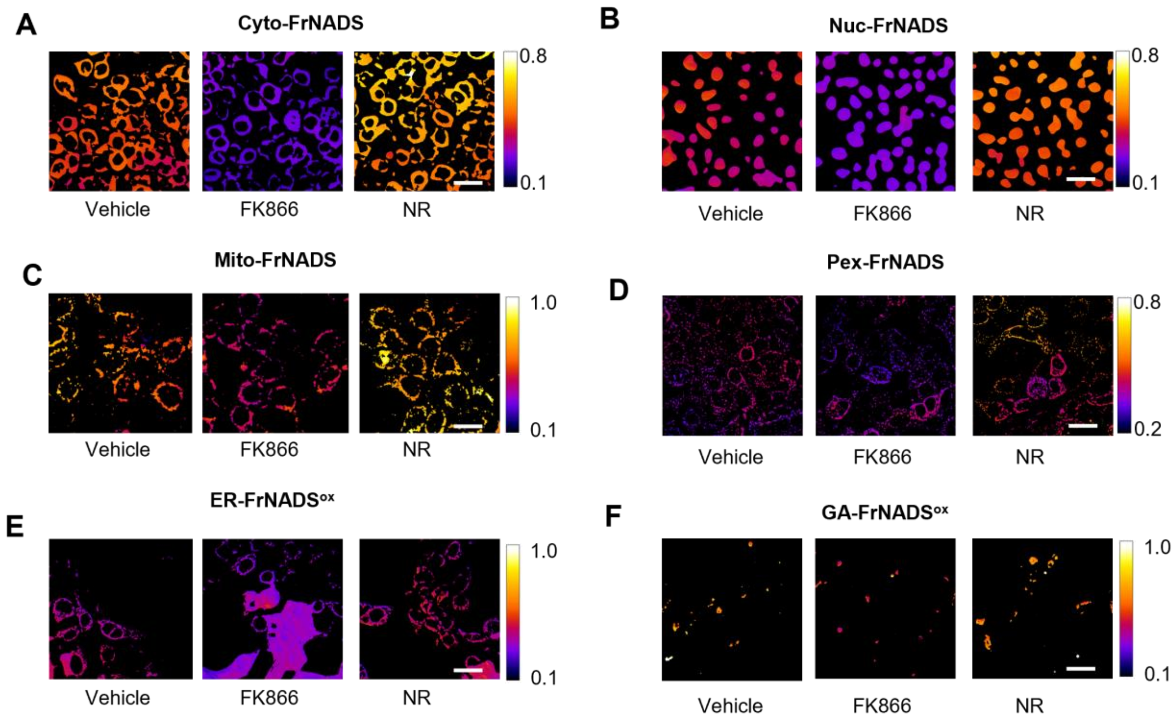


**Figure S3. Representative ratiometric imaging of subcellular NAD$^+$ pools using compartmentalized FrNADS, related to Figure 2.** (A-F) Representative pseudo color emission ratio (R/G) images of HEK 293T cells expressing FrNADS targeted to the cytoplasm (A), nucleus (B), mitochondria (C), peroxisomes (D), endoplasmic reticulum (E, utilizing the ER-FrNADS$^{ox}$ variant), and Golgi apparatus (F, utilizing the GA-FrNADS$^{ox}$ variant). Cells were treated with vehicle (control), the NAMPT inhibitor FK866, or the NAD$^+$ precursor NR for 24 h prior to imaging. Calibration bars showing the dynamic range of the emission ratio are provided on the right of each panel set. These images correspond to the quantitative statistical data presented in Figure 2A. Scale bars, 30 μm.

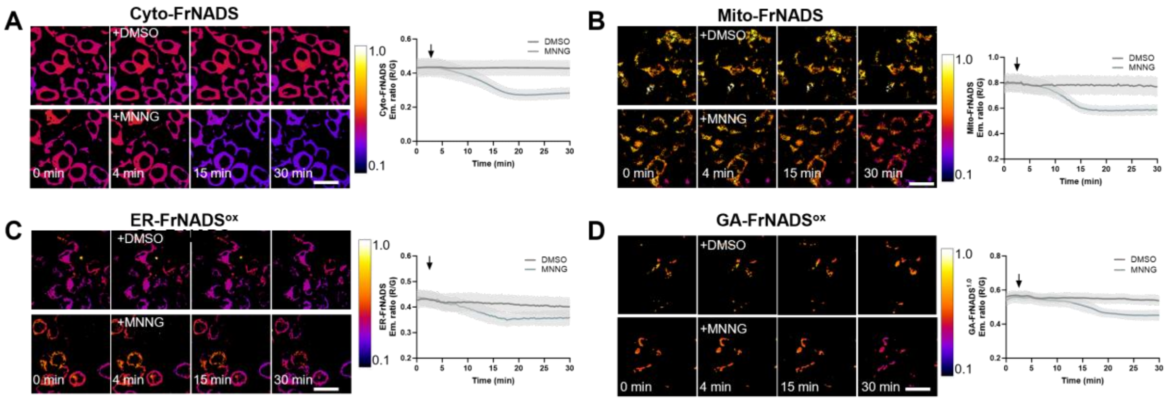


**Figure S4. Real-time monitoring NAD$^+$ depletion across subcellular compartments upon MNNG administration, related to Figure 2.** Representative time-lapse pseudo color emission ratio (R/G) images (left panels) and corresponding quantitative kinetic traces (right panels) of HEK 293T cells expressing FrNADS targeted to the cytoplasm (A), mitochondria (B), endoplasmic reticulum (C), and golgi apparatus (D). Scale bars, 30 μm. Data are presented as mean ± SD. ****, $P < 0.0001$.

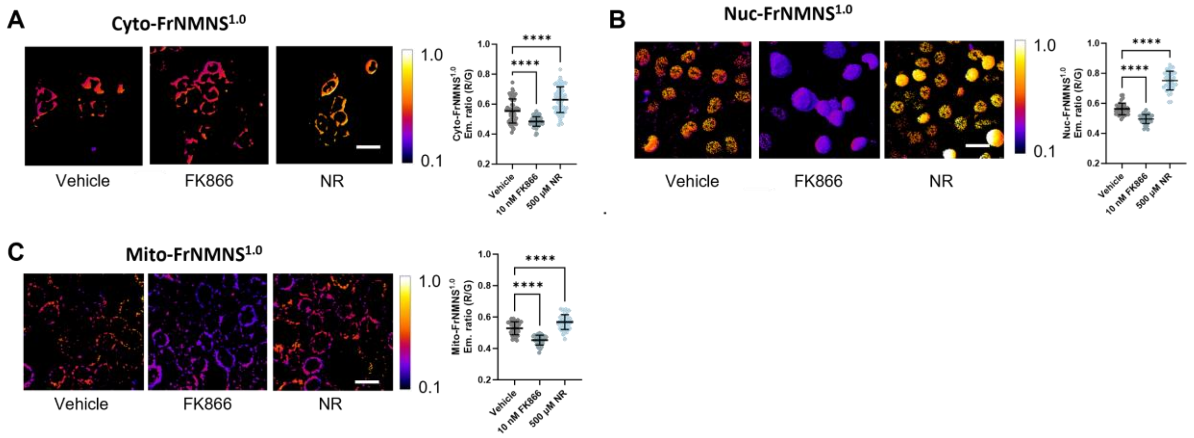


**Figure S5. Validation of compartmentalized FrNMNS$^{1.0}$ biosensors for monitoring subcellular NMN pools.** (A-C) Representative pseudo color emission ratio (R/G) images (left panels) and corresponding quantitative analysis (right panels) of HEK 293T cells expressing FrNMNS$^{1.0}$ targeted to the cytoplasm (A), nucleus (B), and mitochondria (C). To validate the biosensor's responsiveness to NMN fluctuations, cells were treated with vehicle, the NAMPT inhibitor FK866 or precursor NR. Scale bars, 30 μm. Data are presented as mean ± SD, statistical significance was determined using one-way ANOVA analysis followed by Dunnett's multiple comparisons test. ****, $P < 0.0001$.

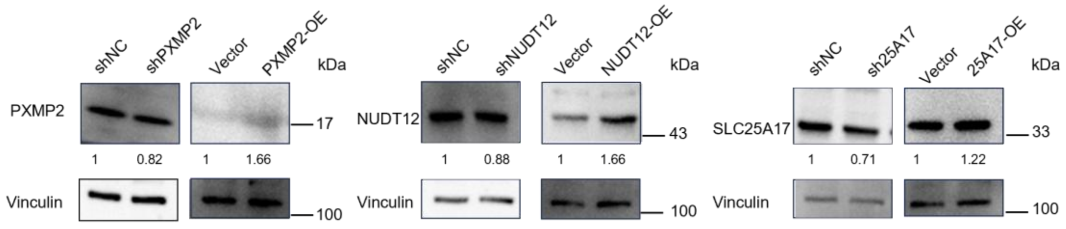


**Figure S6. Validation of genetic perturbation of Pxmp2, Nudt12, and Slc25a17. Related to Figure 3.** Representative immunoblotting of PXMP2, NUDT12, and SLC25A17 in HepG2 cells transduced with shRNA targeting each gene or non-targeting control (shNC). Vinculin was used as loading control.

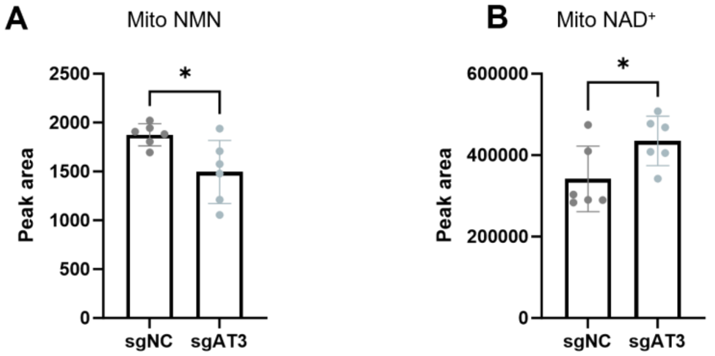


**Figure S7. Related to Figure 4.** Mitochondrial NMN (A) and $NAD^+$ (B) content in NMNAT3 knock down relative to control HepG2 cells determined by LC-MS. Data are presented as mean ± SD, statistical significance was determined using two-tailed Student's t-test. *, $P < 0.05$.

**A**

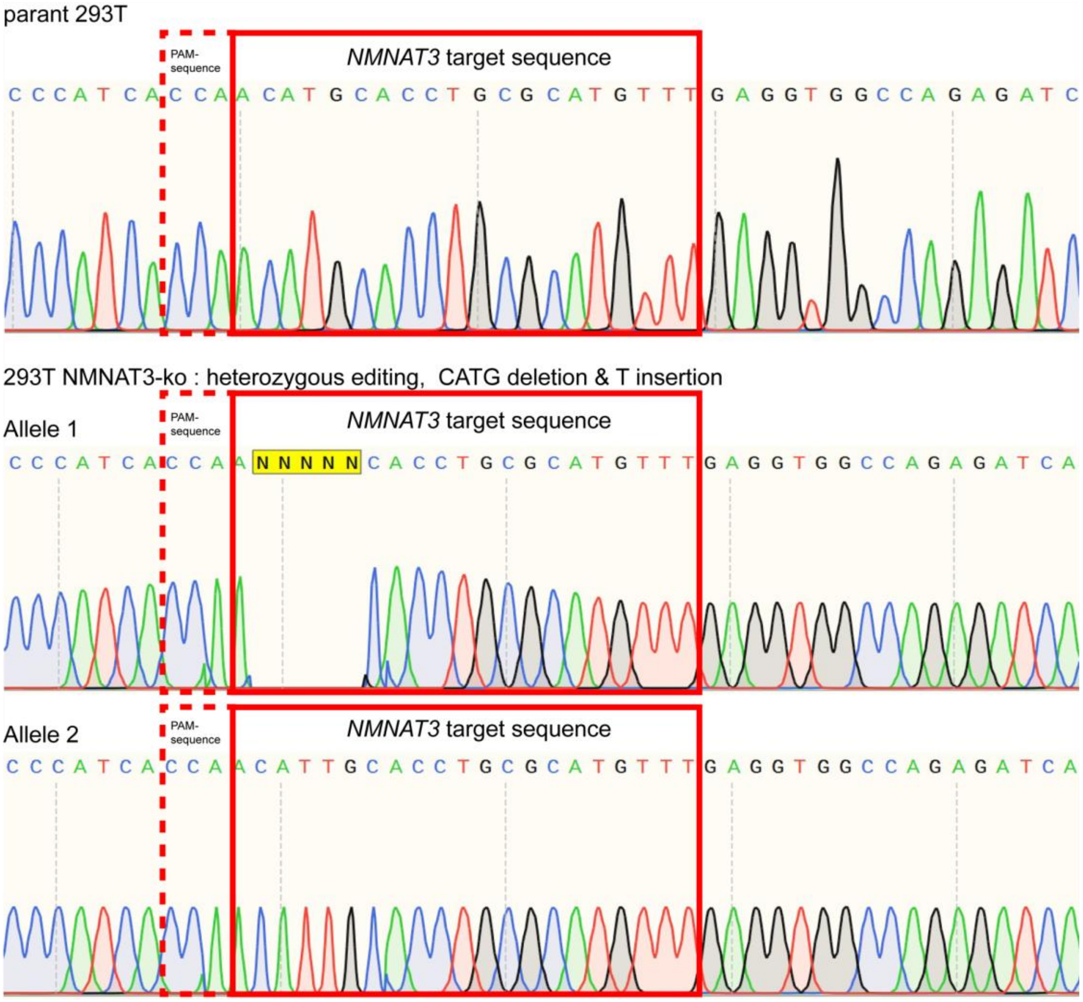


**B**

| sample | pos | ref | alt | dp | dp_alt | freq | dp_alt_fr | var_type | var_seq | indel | seq |
|---|---|---|---|---|---|---|---|---|---|---|---|
| G22606126567-AT3KO | 523-529 | AACATGC | AA-----C | 2229 | 1129 | 0.507 | 555,574 | DEL[2],DEL[2] | CA,TG | -4 | GCTCCTTTAACCCCATCACC[AACA-TGC/AA-----C]ACCTGCGCATGTTTGAGGTG |
| G22606126567-AT3KO | 523-529 | AACATGC | AACATTGC | 2229 | 1100 | 0.494 | 518,582 | INS[1] | T | 1 | GCTCCTTTAACCCCATCACC[AACA-TGC/AACATTGC]ACCTGCGCATGTTTGAGGTG |

**C**

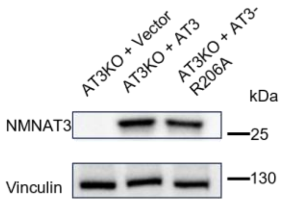


**Figure S8. Validation of HEK 293T *NMNAT3* KO clones.** (A) Nanopore sequencing analysis of the critical region targeted by *NMNAT3* specific sgRNA. (B) Summary of the editing events in the region targeted by *NMNAT3* sgRNA. (C) Immunoblotting analysis of NMNAT3 overexpression in HEK 293T cells.

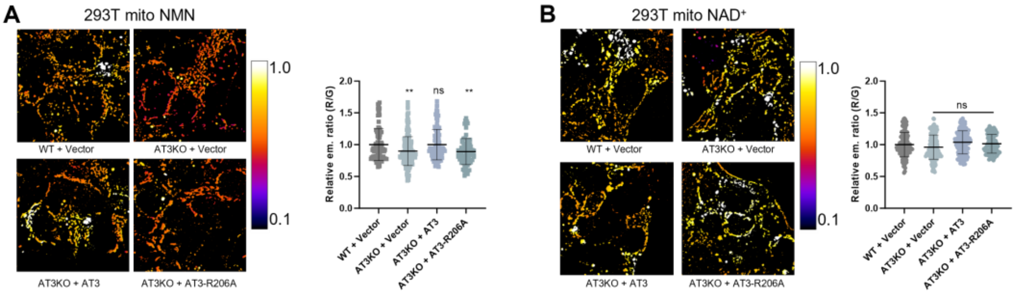


**Figure S9. NMNAT3 function as NAD$^+$ hydrolyzing enzyme in HEK 293T mitochondria.** Representative pseudocolor FRET emission ratio (RFP/GFP) images of HEK 293T cells expressing mitochondrial matrix targeting FrNMNS$^{1.0}$ (A) or FrNADS (B), illustrating the relative levels of mitochondrial NMN and NAD$^+$ under NMNAT3 knockout or overexpression.

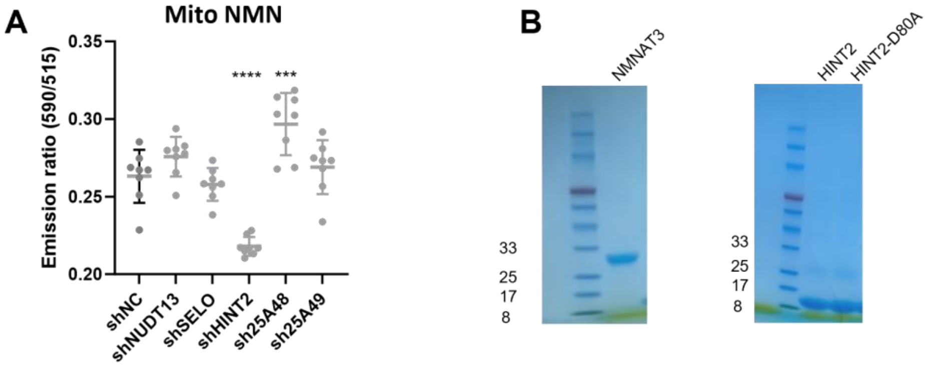


**Figure S10. Related to figure 6**. (A) Scatter plot showing emission ratio (590/515) of the mito-FrNMNS$^{1.0}$ biosensor in HEK 293T expressing shRNA targeting candidate genes. Intensity at 590 nm and 515 nm were obtained from fluorescence microplate reader. (B) SDS-Page gels of purified NMNAT3 and HINT2. Data are presented as mean ± SD of biological replicates, statistical significance was determined using one-way ANOVA analysis followed by Dunnett's multiple comparisons test. ns, $P \geq 0.05$; ***, $P < 0.001$; ****, $P < 0.0001$.

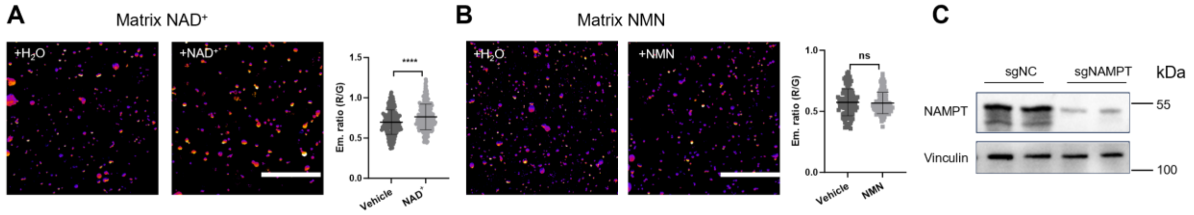


**Figure S11. Related to figure 7**. (A) Representative pseudo color fluorescence images (left) and corresponding quantification (right) of purified mitochondria expressing the matrix-targeted $NAD^+$ biosensor following incubation with vehicle (ultrapure water) or 1 mM $NAD^+$ for 30 min at RT. (B) Representative pseudo color fluorescence images (left) and corresponding quantification (right) of purified mitochondria expressing the matrix-targeted NMN biosensor following incubation with vehicle (ultrapure water) or 100 μM NMN for 30 min at RT. (C) Western blot analysis of *NAMPT* knockdown efficiency. Scale bars, 5 μm. Data are presented as mean ± SD; statistical significance was determined using two-tailed Student's t-test. ns, $P \geq 0.05$; ****, $P < 0.0001$.
